# Cortical ripples mediate top-down suppression of hippocampal reactivation during sleep memory consolidation

**DOI:** 10.1101/2023.12.12.571373

**Authors:** Justin D. Shin, Shantanu P. Jadhav

**Affiliations:** Neuroscience Program, Department of Psychology, and Volen National Center for Complex Systems, Brandeis University, Waltham, MA 02453, USA

## Abstract

Consolidation of initially encoded hippocampal representations in the neocortex through reactivation is crucial for long-term memory formation, and is facilitated by the coordination of hippocampal sharp-wave ripples (SWRs) with cortical oscillations during non-REM sleep. However, the contribution of high-frequency cortical ripples to consolidation is still unclear. We used continuous recordings in the hippocampus and prefrontal cortex (PFC) over the course of spatial learning and show that independent PFC ripples, when dissociated from SWRs, predominantly suppress hippocampal activity in non-REM sleep. PFC ripples paradoxically mediate top-down suppression of hippocampal reactivation, which is inversely related to reactivation strength during coordinated CA1-PFC ripples. Further, we show non-canonical, serial coordination of ripples with cortical slow and spindle oscillations. These results establish a role for cortical ripples in regulating consolidation.

## Main Text

Systems memory consolidation is the process through which newly formed memories are transferred to cortical regions for long term storage, and models of memory consolidation such as the standard systems theory of consolidation and multiple trace theory posit a predominantly unidirectional interaction between the hippocampus and cortical regions (*1-3*). This consolidation process is facilitated through the coordination of patterns of activity that span a range of timescales and oscillatory frequencies and has been extensively studied in the context of the hippocampal-prefrontal cortical circuit. The most prominent of these activity patterns are hippocampal sharp-wave ripples (SWRs), a phenomenon critical for memory consolidation (*4*). These high frequency bursts during non-REM (NREM) sleep reactivate hippocampal representations of previous experiences, engage the prefrontal cortex (PFC), and are associated with brain wide changes in activity, hypothesized to support integration of mnemonic representations in distributed cortical circuits (*5-8*). Similarly, high frequency ripple bursts have been observed in the neocortex, including PFC, that synchronize local activity and have been reported to be coupled to hippocampal SWRs (*9, 10*). However, whether and how PFC ripples reciprocally engage hippocampal activity, and especially hippocampal reactivation, in support of consolidation is unclear.

### Independent and coordinated ripple events across CA1 and PFC

Here, we simultaneously recorded populations of CA1 (n = 3,316) and PFC (n = 2,474) neurons (fig. S1) from animals (n = 8 rats) in NREM sleep (mean = 13.24 ± 0.83 min per epoch, fig. S2) throughout the course of learning a spatial alternation task that requires hippocampal-prefrontal interactions (*11, 12*). Activity was monitored across multiple run and sleep sessions during learning. We detected cortical ripples in PFC as previously described (*9, 10*), and classified them as either temporally coupled to hippocampal SWRs (i.e., coordinated ripples) or independent (Fig. 1A-D). We thus separated ripples based on the interregional overlap of these events as independent PFC ripples, independent CA1 SWRs (termed independent SWRs), and coordinated CA1-PFC ripples (fig. S3,S4). Independent PFC ripple and independent CA1 SWR events were more prevalent than coupled CA1-PFC ripples events in NREM sleep (fig. S3B; independent PFC ripple rate: 0.46 ± 0.02 Hz; independent CA1 ripple rate: 0.67 ± 0.02 Hz; coordinated ripple rate: 0.16 ± 0.01 Hz). We hypothesized that these two kinds of events, independent and coordinated, may subserve different mnemonic processes within the hippocampal-prefrontal network, and found key differences. Coordinated events were longer in duration and had higher cell participation (fig. S5A,C), consistent with previous reports that long duration events are important for performing behavioral tasks with a high memory demand (*13*). The rate of independent PFC ripples increased over time and learning as compared to CA1 SWRs, and was correlated with task performance in the subsequent behavioral session (Fig. 1E, fig. S3C). Conversely, *within* each individual sleep session, independent SWR rate decreased over time, shifting the balance between independent and coordinated SWRs over the course of a sleep session (fig. S3D,E). The rate of coordinated CA1-PFC ripples also increased with learning and was correlated with the increase in overall PFC ripple rate, suggesting a role for PFC in driving the increase in coupling during later sleep epochs (Fig. 1F, fig. S3F). Finally, coordinated CA1-PFC ripples were also associated with higher spindle power (Fig. 1G), indicating enhanced cross-frequency coupling between the hippocampus and PFC during coordination, which is critical for memory consolidation (*14-16*).

**Fig. 1.**
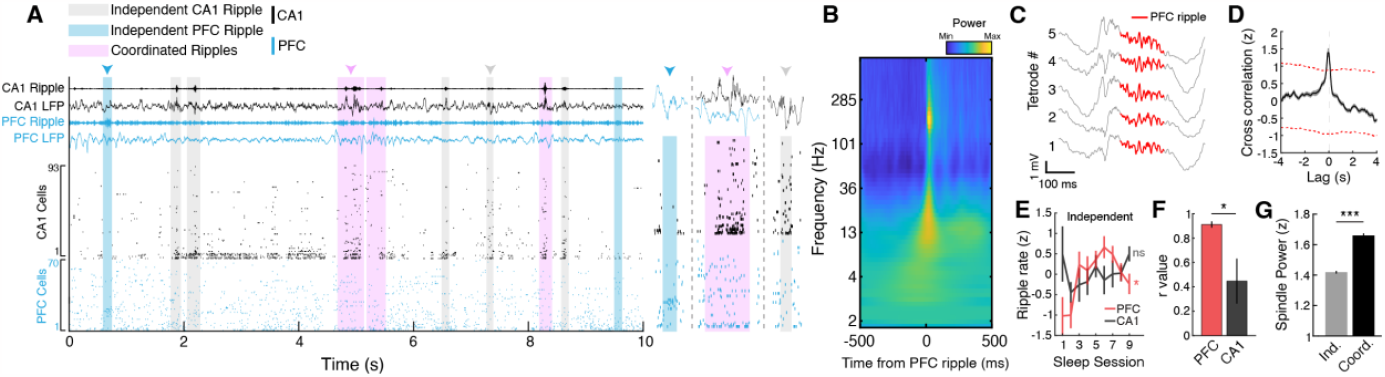
Independent and coordinated ripple events in CA1 and PFC. (**A**) Example CA1 and PFC raster plot illustrating the occurrence of independent CA1 and PFC ripples and coordinated events. Events indicated by the arrowheads are expanded on right to show LFP and single unit activity. (**B**) PFC ripple triggered spectrogram combined across all animals (n = 8 rats) and epochs (n = 72 epochs; n = 37,335 ripples). Note the associated spindle band activity (12-16 Hz) during these events. (**C**) Example PFC ripple event shown across 5 tetrodes. Detected ripple event is highlighted in red. (**D**) Cross-correlation between all ripple events in CA1 and PFC. Red dashed line indicate the 95% confidence intervals for jittered data. (**E**) Independent ripple rate over the 9 sleep epochs (left, PFC: F = 2.44, *p = 0.03; CA1: F = 1.18, p = 0.34, main effect of time, repeated measures ANOVA). (**F**) The correlations between overall and coordinated ripple rate over time for CA1 and PFC events suggest PFC driven coupling (PFC, r = 0.91 ± 0.03; CA1, r = 0.45 ± 0.18, *p = 0.01). (**G**) Peak spindle power during independent vs coordinated PFC ripple events (Independent = 1.42 ± 0.01, Coordinated = 1.66 ± 0.01, ***p = 2.11×10^−66^).

### Bidirectional modulation of CA1 neurons during PFC ripples

Previous reports have shown that PFC neurons are bidirectionally modulated by hippocampal SWRs (*17-19*), so here we examined the effect of independent PFC ripples on CA1 neuronal activity. Interestingly, a large majority of CA1 cells, both putative pyramidal cells and interneurons (classification shown in fig. S6), were suppressed during independent PFC ripples, while PFC activity was enhanced by the local ripples (Fig. 2A,B; interneuron modulation shown in figs. S7A). In addition, suppression in CA1 during PFC ripples peaked prior to PFC excitation (Fig. 2C), indicating attenuation of CA1 activity through a cortical-hippocampal-cortical loop that may enhance information transfer within the neocortex (*20, 21*). The neuronal activity modulation in CA1 and PFC was dependent on the ripples examined (Fig. 2A, fig. S8A-C), either local ripples in CA1/ PFC or coordinated ripples in CA1-PFC, especially with coordinated ripples predominantly driving excitation in both regions in contrast to independent cortical ripples, highlighting that these distinct events may differ in their contribution to mnemonic processes.

**Fig. 2.**
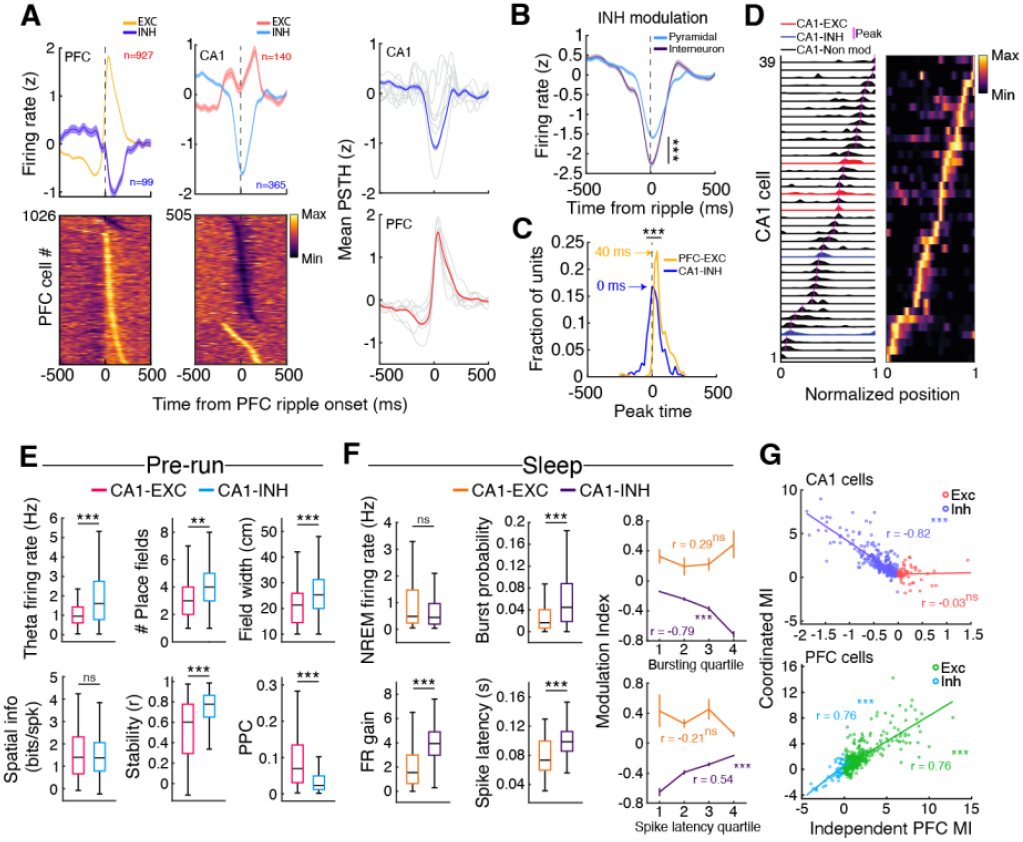
Predominant suppression of CA1 activity during independent PFC ripples. (**A**) Independent PFC ripple triggered modulation in PFC (left) and CA1 (middle). (Right) Overall modulation in each area showing neurons excited or suppressed during ripples, with predominantly excitation in PFC and predominantly suppression in CA1. Gray lines indicate individual animals. (**B**) The degree of suppression was higher in INH interneurons as compared to pyramidal cells (±200 around ripple onset; \*\**\**p = 1.37×10^−9^). (**C**) Timing of peak suppression for CA1 INH cells (interneurons and pyramidal cells) and PFC EXC cells (Comparison of peak timing distributions, ***p = 4.90×10^−46^). (**D**) Example firing maps for CA1 EXC, INH, and non-modulated cells during one trajectory. (**E**) Differences in spatial properties for CA1 EXC and INH pyramidal cells during the run session preceding the sleep epoch in which the cells were modulated (Theta firing rate, EXC = 1.20 ± 0.10 Hz, INH = 2.05 ± 0.10 Hz, ***p = 2.08×10^−6^; Number of fields, EXC = 3.3 ± 0.17, INH = 3.99 ± 0.11, **p = 0.002; Field width, EXC = 21.67 ± 0.91 cm, INH = 26.56 ± 0.61 cm, ***p = 1.1×10^−4^; Spatial information (bits/spike), EXC = 1.52 ± 0.12 bits, INH = 1.49 ± 0.06 bits, p = 0.83; Stability, EXC = 0.54 ± 0.03, INH = 0.72 ± 0.01, ***p = 9.01×10^−7^; Pairwise phase consistency, EXC = 0.10 ± 0.01, INH = 0.05 ± 0.004, ***p = 3.31×10^−10^). (**F**) (Left) Mean NREM firing rate and intra-SWR firing dynamics for CA1 EXC and INH cells during coordinated ripples (NREM firing rate, EXC = 1.07 ± 0.11 Hz, INH = 0.81 ± 0.05 Hz, p = 0.18; FR gain, EXC = 2.02 ± 0.16, INH = 4.07 ± 0.10, ***p = 2.88×10^−27^; Burst probability, EXC = 0.031 ± 0.006, INH = 0.075 ± 0.005, ***p = 6.38×10^−5^; Spike latency, EXC = 0.080 ± 0.006, INH = 0.099 ± 0.001, ***p = 1.61×10^−5^). (Right) Relationship between SWR bursting incidence (top, EXC: r = 0.29, p = 0.08, INH: r = -0.79, ***p = 5.20×10^−69^) or spike latency for CA1 EXC and INH cells (bottom, EXC: r = -0.21, p = 0.21, INH: r = 0.54, *** p = 9.51×10^−25^) with modulation index (MI) during independent PFC ripple events. (**G**) Relationship between MI during coordinated ripples and independent PFC ripples (top, CA1, EXC: r = 0.03, p = 0.82; INH: r = -0.82, ***p = 6.84×10^−80^; bottom, PFC, EXC: r = 0.76, ***p = 1.34×10^−92^; INH: r = 0.76, ***p = 1.51×10^−10^). Note the negative relationship for CA1 INH cells, indicating that stronger the recruitment during coordinated ripples, the more inhibited the cells are during independent PFC ripples.

Further examination of the inhibited (INH, n = 365) and excited (EXC, n = 140) CA1 pyramidal neurons during run sessions revealed representational differences (Fig. 2D,E). INH cells had higher theta firing rates, had wider and more numerous fields, and had greater stability between sessions, whereas EXC cells were more strongly locked to the hippocampal theta oscillation (Fig. 2E, fig. S8D). During NREM sleep, EXC and INH cells had similar baseline firing rates but had distinct intra-SWR firing dynamics. EXC cells were generally active earlier in SWR events and INH cells were more strongly excited (Fig. 2F). The degree of CA1 modulation by PFC ripples was also predictive of intra-CA1-SWR dynamics (Fig. 2F, right) and was negatively correlated with modulation during coordinated CA1-PFC ripples (Fig. 2E), indicating a role for PFC ripples in influencing the content of hippocampal activity during coordinated ripples. In particular, the suppression of CA1 INH cells by PFC ripples that are strongly recruited during coordinated CA1-PFC ripples suggests that independent PFC ripples suppress hippocampal reactivation during spatial learning. Similar relationships were also found in the population of putative CA1 interneurons, demonstrating a broad top-down influence of PFC ripples on CA1 activity (fig. S7B-D). Thus, PFC ripples delineate transient intervals where hippocampal activity is largely suppressed, potentially modifying circuit dynamics for effective information processing or transfer.

### Top-down suppression of CA1 reactivation

We then examined population activity in CA1 and PFC during these ripple events using assembly analysis to explicitly look at reactivation during ripples (Fig. 3A,C). Corroborating the single unit results, assemblies within a region (CA1 n = 585, PFC n = 320) were robustly reactivated during their respective local ripple events (Fig. 3B,D, fig. S9B,C) and were strongly co-active during coordinated CA1-PFC ripple events as expected for a memory consolidation process (fig. S9D,G). CA1 EXC cells had higher weights and contribution to reactivation strength (Fig. 3D, fig. S9I), possibly owing to the stronger theta phase locking within this population during run sessions (Fig. 2E, fig. S8D). Assemblies in CA1 and PFC exhibited gradients of reactivation strengths across events (fig. S10), and the degree of reactivation strength in CA1 was negatively correlated across independent PFC ripples and coordinated CA1 SWRs (Fig. 3E), confirming that assemblies that are strongly reactivated during coordinated CA1-PFC ripples are highly suppressed by independent PFC ripples. This CA1 suppression was also strongest during the later sleep sessions when PFC ripple rate peaked (Fig. 1E, fig. S9E).

**Fig. 3.**
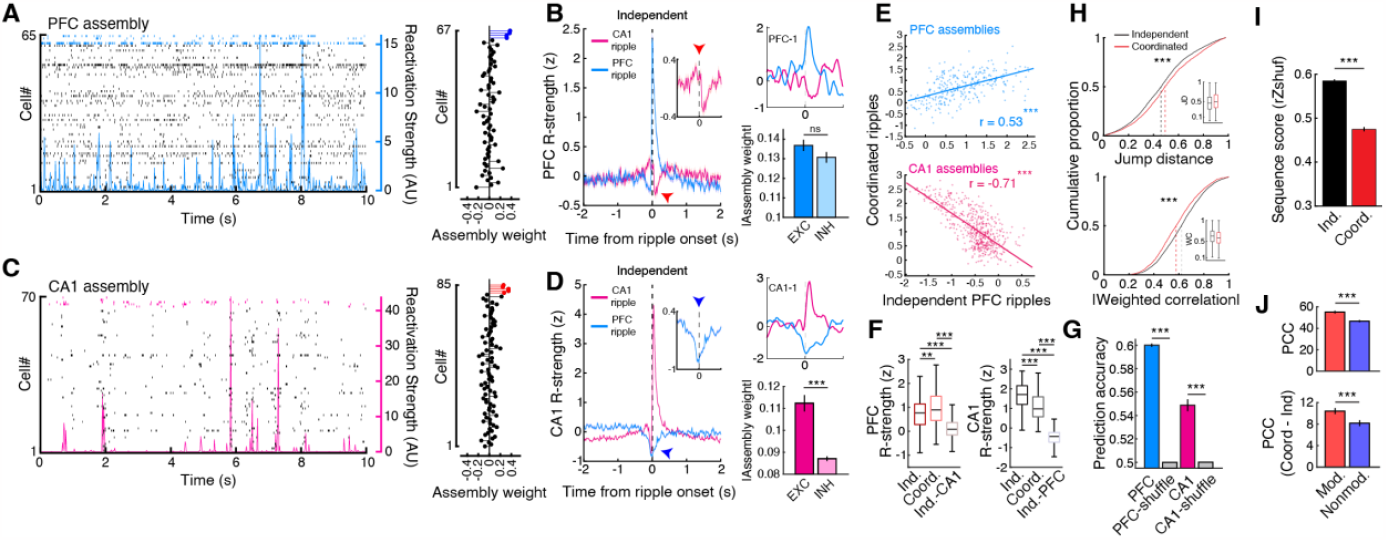
Assembly reactivation during independent and coordinated ripple events. (**A**) (Left) Example PFC raster plot during NREM sleep with the reactivation strength of an assembly overlaid. (Right) Neuron weights for the plotted assembly on the left, with member cells highlighted in blue. (**B**) (Left) Independent ripple triggered PFC reactivation strength. (Right) Example PFC assembly aligned to ripple events (top) and the absolute weights of EXC and INH PFC cells (modulation during independent CA1 SWRs, EXC = 0.137 ± 0.003, INH = 0.131 ± 0.003, p = 0.37). (**C**) and (**D**) Same as in (**A**) and (**B**) but for CA1 (modulation during independent PFC ripples, EXC = 0.112 ± 0.004, INH = 0.087 ± 0.001, ***p = 8.04×10^−7^). (**E**) Relationship between Z-scored assembly reactivation strength during independent PFC ripples and coordinated ripples (PFC: r = 0.53, ***p = 1.28×10^−23^; CA1: r = -0.71, ***p = 4.25×10^−87^). Note that PFC reactivation strength is correlated across events, whereas CA1 assemblies that are highly suppressed during independent PFC ripples are strongly reactivated during coordinated ripples. (**F**) Reactivation strength during within-area independent and coordinated ripples and during opposing area independent events (PFC: Independent-PFC = 0.71 ± 0.03, Coordinated-PFC = 0.96 ± 0.04, Independent-CA1 = 0.13 ± 0.02, **p = 0.0023, ***p = 3.21×10^−33^, ***p = 2.12×10^−53^ for independent-PFC vs coordinated-PFC, independent-PFC vs independent-CA1, and coordinated-PFC vs independent-CA1, respectively; CA1: Independent-CA1 = 1.67 ± 0.03, Coordinated-CA1 = 1.11 ± 0.03, Independent-PFC = -0.51 ± 0.02, ***p = 2.44×10^−18^, ***p = 7.45×10^−239^, ***p = 1.14×10^−128^ for independent-CA1 vs coordinated-CA1, independent-CA1 vs independent-PFC, and coordinated-CA1 vs independent-PFC, respectively; Kruskal-Wallis test, Bonferroni corrected for multiple comparison). (**G**) Prediction of intra-area ripple type (independent or coordinated) using spike count data (PFC: data = 0.6 ± 0.001, shuffle = 0.5 ± 4.64×10^−5^, ***p = 3.39×10^−50^; CA1: data = 0.55 ± 0.005, shuffle = 0.50 ± 1.37×10^−4^, ***p = 1.28×10^−26^). (**H**) Jump distance and absolute weighted correlation for significant CA1 replay events decoded during independent CA1 SWRs and coordinated ripples (Jump distance: Independent = 0.468 ± 0.003, Coordinated = 0.505 ± 0.005, ***p = 2.73×10^−11^; Weighted correlation: Independent = 0.620 ± 0.002, Coordinated = 0.585 ± 0.003, ***p = 1.10×10^−17^). (**I**) Sequence degradation (rZ_shuf_) of independent and coordinated replay events after shuffling individual cells’ linearized firing fields (Independent = 0.585 ± 0.004, Coordinated = 0.475 ± 0.005, ***p = 5.46×10^−76^). (**J**) Per-cell contribution (PCC) to all significant replay for CA1 modulated vs non-modulated cells (top, modulated = 55.04 ± 1.28, non-modulated = 46.65 ± 1.02, ***p = 2.39×10^−7^) and the difference in PCC (coordinated minus independent) to independent and coordinated events (bottom, modulated = 10.44 ± 0.57, non-modulated = 8.17 ± 0.55, ***p = 9.88×10^−4^).

Interestingly, the degree of suppression of CA1 assemblies during independent PFC ripples was related to assembly reinstatement in subsequent behavioral sessions (fig. S9H), suggesting a role for this top-down suppression in modifying subsequent CA1 coactivity patterns. In addition, both reactivation strength and within-area coactivity in PFC and CA1 differed across ripple events, and population activity in PFC and CA1 was able to distinguish between independent and coordinated ripples in each respective area (Fig. 3F,G, fig. S9F). We observed similar relationships for joint CA1-PFC reactivation strength. Joint CA1-PFC coactivity differed between independent and coordinated SWRs, and showed a similar negative relationship between reactivation strength during coordinated ripples vs. independent PFC ripples (fig. S11). These results indicate that independent and coordinated ripples underlie different modes of information exchange between CA1 and PFC, and further that stronger the reactivation in CA1 during coordinated CA1-PFC ripples, the stronger the suppression during PFC ripples.

Since it has been previously demonstrated that hippocampal reactivation and replay are separable components of the dynamic hippocampal code (*22*), we examined how association with PFC ripples may impact CA1 replay. We found that replay events associated with coordinated SWRs were less sequential in structure, having higher normalized jump distances and lower weighted correlations than independent-SWR associated replay (Fig. 3H, fig. S12). Sequential unit spiking was also more consistent, as measured by rank-order correlation (*23*), and had higher dimensionality during independent SWRs, further highlighting the influence of PFC on SWR content (fig. S12D,E). Coordinated-SWR replay exhibited a larger degree of sequence degradation when place cell templates are shuffled as compared to independent-SWR replay, possibly due to a decrease in overall reactivation strength and coactivity during these events (Fig. 3F,I, fig. S9F). Furthermore, PFC ripple modulated CA1 cells had higher per-cell contribution (PCC) to replay than non-modulated cells (Fig. 3J, top). Although both modulated and non-modulated cells contributed more strongly to coordinated-SWR replay, which is expected due to the greater sequence degradation observed in these events, modulated cells had higher PCC differences (Fig. 3J, bottom). This overall higher contribution of modulated cells to coordinated-SWR replay and the differences observed between independent and coordinated events further suggest a role for PFC in shaping the content of hippocampal reactivation and replay during sleep.

### Sequential coupling of sleep oscillations across CA1 and PFC

In addition to the recent observation of temporal coupling of hippocampal-cortical ripples contributing to memory consolidation, the coordination of hippocampal SWRs with cortical slow oscillations (SOs) and spindles is known to be important for consolidation (*14, 24, 25*). To this end, we investigated the incidence of ripple events in the context of these oscillations that span multiple timescales. Independent and coordinated CA1 SWRs exhibited differential coupling with spindles and SOs, with only coordinated CA1-PFC ripples associated with cortical spindles and not independent CA1 SWRs (Fig. 4A, fig. S13). Population activity in CA1 and PFC also had unique responses surrounding independent vs. coordinated events (fig. S14). Additionally, CA1 replay events, specifically during coordinated CA1-PFC ripples, were enriched during up states of the SO (Fig. 4B,C, fig. S15B). These results, along with the finding that coordinated CA1-PFC ripples were associated with a higher cortical spindle power (Fig. 1G), led us to hypothesize distinct temporal organization of independent and coordinated ripple events with slow oscillations and spindles.

**Fig. 4.**
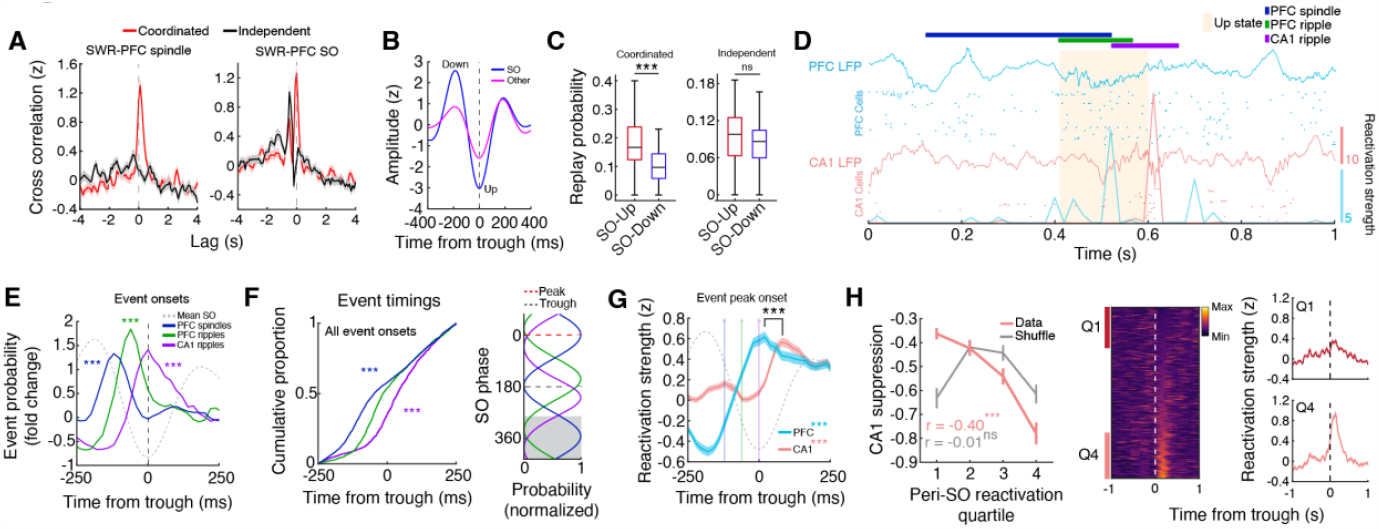
Serially coupled spindles, ripples, and slow oscillations. (**A**) Cross-correlation between independent or coordinated CA1 SWRs with PFC spindles and SOs (trough time). Note that only coordinated CA1-PFC ripples tend to be associated with spindle oscillations. Furthermore, the timing of SWRs relative to SOs is different between these events, with coordinated ripples occurring primarily at the trough and independent SWRs preceding SO troughs. (**B**) Average waveforms for SOs and other, lower amplitude slow waves extracted from PFC LFPs. (**C**) Probability of coordinated and independent ripple associated CA1 replay during up and down states (Coordinated: up = 0.19 ± 0.02, down = 0.11 ± 0.01, ***p = 2.30×10^−4^; Independent: up = 0.09 ± 0.01, down = 0.08 ± 0.01, p = 0.18). (**D**) Raster showing the reactivation strengths of example CA1 and PFC assemblies in relation to LFP events. Note the sequential organization of events with PFC leading CA1. (**E**) Event probability fold change of spindles and ripples aligned to SO troughs (Spindles, ***p = 2.84×10^−14^; PFC ripples, ***p = 1.66×10^−7^; CA1 ripples, ***p = 3.86×10^−5^, ttest vs 0), showing the spindle-PFC ripple-CA1 ripple timing. (**F**) (Left) Cumulative proportion of event timings within ±250 ms of SO troughs (Spindles vs PFC ripples, ***p = 2.41×10^−16^, PFC ripples vs CA1 ripples, ***p = 4.62×10^−13^), and (Right) preferred SO phase for each event type (Spindles vs PFC ripples, U^2^ = 2.63, ***p = 5.73×10^−23^, PFC ripples vs CA1 ripples, U^2^ = 2.54, ***p = 3.61×10^−22^, Watson’s U^2^ test). Gray shaded area indicates repeated data for visualization. (**G**) SO trough aligned CA1 and PFC reactivation strength during SO associated coordinated events where PFC ripples precede CA1 SWRs (PFC vs shuffle, ***p = 4.91×10^−19^; CA1 vs shuffle, ***p = 2.12×10^−39^; PFC vs CA1 timing, ***p = 7.96×10^−5^). (**H**) (Left) Relationship between independent PFC ripple associated CA1 suppression and SO upstate CA1 reactivation strength (data: r = -0.40, ***p = 3.01×10^−21^, shuffle: r = -0.01, p = 0.86), again showing a negative relationship. (Middle) SO trough aligned reactivation strength for all CA1 assemblies. (Right) Z-scored SO trough aligned CA1 reactivation strength plotted for the first and fourth quartiles.

We observed an overall higher prevalence of coordinated events where PFC ripples precede CA1 SWRs (fig. S16A), which is consistent with PFC ripples potentially driving ripple coordination (Fig. 1F), and therefore focused on these events for the subsequent analyses to investigate sequential coupling of events in CA1 and PFC. We quantified the onset of PFC spindles, ripples, and CA1 SWRs relative to the trough (up state) of SOs and calculated the fold change probability of each event around SOs. We found a sequential organization of these oscillatory events that suggests a thalamocortical-hippocampal directionality, with spindles preceding coordinated PFC-CA1 ripple events (Fig. 4D,E) as well as independent PFC ripples (fig. S15D). This temporal relationship with spindles was not present for independent CA1 SWRs (fig. S15D) and differed for coordinated events where CA1 SWRs preceded PFC ripples (fig. S15E). The embedding of PFC ripple events between spindles and CA1 SWRs (Fig. 4E,F) would allow for the coordination of PFC and CA1 assembly reactivation. Indeed, PFC and CA1 reactivation was temporally organized surrounding SO troughs in a manner that is consistent with cortico-hippocampal directionality (Fig. 4G). In addition, the strength of up state aligned reactivation in CA1 assemblies was related to the degree of suppression during independent PFC ripples (Fig. 4H), suggesting a role of PFC ripples in prioritizing certain CA1 memory traces for long-term storage or modification. Additionally, differences in coordinated events (PFC leading vs lagging), such as the timing and content of CA1 and PFC reactivation (fig. S16), suggests a bidirectional coordinated ripple-mediated interaction with potentially distinct roles in consolidation.

Memory consolidation is a dynamic process that is dependent on the interaction between multiple brain systems, and previous studies have shown that enhanced cross-frequency coupling between the hippocampus and PFC enhances performance on memory related tasks (*14, 26*). Our finding that independent and coordinated events across the hippocampal-prefrontal circuit elicit differential responses suggests separate roles in systems memory consolidation. Notably, the suppression of CA1 reactivation by independent PFC ripples is unexpected based on current models of systems consolidation (*1-3*). While coordinated ripple events across hippocampal-neocortical circuits are thought to support memory consolidation through the coordinated reactivation of behaviorally relevant representations (*9, 19*), the roles of independent events in CA1 and PFC are unclear. One possibility is that independent CA1 SWRs (independent meaning uncoupled from PFC ripple activity, specifically) preferentially engage other brain areas to support different aspects of consolidation depending on the nature of the experience. Indeed, there are widespread, bidirectional changes in activity surrounding SWRs (*5, 27*), which could reflect distinct processes if examined on an event-by-event basis. Future studies investigating multiple regions of interest simultaneously will allow for the discretization of events spanning a wide range of brain areas and may reveal a multiplex hierarchy, within which specific components can be selectively engaged depending on mnemonic demands (*28*).

The broad suppression that spans multiple cell types in CA1 during PFC ripples indicates that these cortical ripples have a unique contribution to the consolidation of recently acquired memories. The robust and predictive nature of this suppression suggests that PFC may have a role in modulating or biasing the content of hippocampal reactivation to influence what information to consolidate within the hippocampus, and in turn, the neocortex. Furthermore, modulation during PFC ripples may indicate a mechanism through which the E-I balance within the CA1 circuit is modified to maintain a specific level of criticality (*29*) or to control the excitatory drive in the population of CA1 pyramidal cells, thus increasing the signal to noise ratio to allow for more robust information transfer (*30*). Concurrently, CA1 suppression may also be a mechanism through which memory interference in PFC is attenuated to allow for restricted ripple mediated information transfer and plasticity throughout the neocortex to support memory integration through synchrony for semantic memories (*20, 21, 31*). We hypothesize that PFC ripples may be associated with consolidation within cortical networks while suppressing hippocampal reactivation, and elucidation of the brain-wide processes associated with this suppression will require monitoring multiple cortical areas in future studies. However, it is important to note—given that the suppression of CA1 initiates prior to PFC reactivation during independent PFC ripple events, there are likely other brain regions contributing to this process. Future studies investigating this bidirectional modulation of CA1 in the context of the laminar distribution of cells (deep vs superficial), their projection profiles to different cortical targets, such as PFC and entorhinal cortex, and the state of plasticity (plastic vs rigid) will yield a clearer picture of the multimodal dynamics present in the hippocampus (*32-34*).

The finding that highly suppressed CA1 assemblies are strongly reactivated during coordinated events, which are coupled with spindles and SOs, a salient signature of consolidation, also suggests a mechanism through which PFC prioritizes specific CA1 assemblies for consolidation based on future utility or generalizability (*35*). In the context of memory reconsolidation, suppression may be involved in preventing destabilization of certain salient CA1 memory traces

(*36*). Many studies of reconsolidation utilize experimental paradigms that are performed over the course of multiple days. It has been shown that specific changes in hippocampal dynamics potentially linked to consolidation, namely representational drift (*37*), are a function of experience (*38, 39*); however, our longitudinal study design wherein animals are exposed to multiple behavioral sessions in a day can maintain a specific level of lability in CA1 that is consistent with circuit dynamics underlying reconsolidation. Given the persistent reactivation that occurs in CA1 subsequent to novel experience (*40*), PFC mediated suppression may be a mechanism through which CA1 activity is normalized to a level that is optimal for the induction of transcriptional processes that underlie memory maintenance or integration (*36*).

Previous human fMRI studies have reported hippocampal suppression primarily during retrieval stopping, which is typically investigated in contexts where intrusive memories are proactively suppressed (*41, 42*). Our finding that coordinated SWR associated replay events have lower dimensionality and sequential structure raises the alternative possibility that PFC may be involved in a process that prevents certain neural patterns from being consolidated through disruption of hippocampal sequential activity. SWRs are thought to facilitate retrieval through the reactivation of hippocampal ensembles, but the mechanisms through which these activity patterns are distinguished based on utility for consolidation are unclear. There must be a way for the brain to separate activity that is elicited by actual experience from novel, unrelated activity patterns that are a result of internally generated mechanisms to maintain a faithful rendering of reality (*8, 43*). Although these coordinated SWR associated replay events quantitatively represent sequences of behaviorally relevant activity, the up and downstream brain areas conveying and receiving information respectively may play roles in determining the nature of the top-down influence of PFC on hippocampal sequential activity (*28*). This type of reality monitoring is crucial for the purported role of SWRs in imagination (*8*), and further elucidation of the mechanisms underlying this process may lead to a deeper understanding of how it goes awry in neurological disorders such as schizophrenia (*43*).

Our results highlight the dynamic and multidimensional nature of memory consolidation and uncover a previously unknown PFC ripple mediated phenomenon with a distinct suppressive reactivation role that is involved in modifying hippocampal representations and biasing CA1 reactivation in sleep for effective future behavior. Notably, the ability to distinguish SWR events based on long-range interregional coordination affords the opportunity to study this widespread hippocampal phenomenon in greater detail and may be able to reconcile how SWR diversity may play a role in different aspects of cognition (*33, 44-46*).

## Author contributions

Conceptualization: JDS, SPJ

Methodology: JDS

Investigation: JDS

Visualization: JDS

Funding acquisition: JDS, SPJ

Supervision: SPJ

Writing – original draft: JDS

Writing – review & editing: JDS, SPJ

## Data and materials availability

Data underlying these results will be uploaded in the NWB (Neurodata Without Borders) format to DANDI (ID#TBD) upon acceptance. Code to replicate these results will be available on our lab GitHub (http://github.com/JadhavLab/PrefrontalRipples) upon acceptance.

## Supplementary Materials

Materials and Methods

Figs. S1 to S16

## Supplementary Materials for

## Materials and Methods

### Experimental model and subject details

All experimental procedures were approved by the Institutional Animal Care and Use Committee at Brandeis University and conformed to US National Institutes of Health guidelines. Eight adult male Long-Evans rats (450-600 g, 4-6 months; RRID: RGD_2308852) were used in this study. Animals were individually housed and kept on a 12-hr regular light/dark cycle.

### Behavior

Animals were trained on a novel W-maze in a single day, as previously described (*47*). Briefly, during the experimental day, all animals ran eight 15-20 minute behavioral session on a W-maze interleaved with nine 20-30 minute rest sessions in a sleep box. Reward (evaporated milk) was automatically delivered at each of the 3 reward wells upon completion of a correct trajectory. At the end of a behavioral session, animals were transferred to an opaque sleep box to rest.

### Surgical implant and electrophysiology

Surgical implantation procedures were as previously described (*47*). Animals (n = 8) were implanted with a microdrive array containing 30-64 independently moveable tetrodes targeting right dorsal hippocampal region CA1 (−3.6 mm AP and 2.2 mm ML) and right PFC (+3.0 mm AP and 0.7 mm ML) (fig. S1). For 64 tetrode recordings (n = 1 animal), CA1 and PFC were targeted bilaterally. On the days following surgery, hippocampal tetrodes were gradually advanced to the desired depths using characteristic EEG patterns (sharp wave polarity, theta modulation) and neural firing patterns as previously described (*47*). A CA1 tetrode in corpus callosum served as hippocampal reference, and another tetrode in overlying cortical regions with no spiking signal served as prefrontal reference. A ground (GND) screw installed in the skull overlying cerebellum also served as a reference. All spiking activity and ripple-filtered LFPs (150-250 Hz; CA1 and PFC; see below) were recorded relative to the local reference tetrode. For detection of slower frequency components (i.e., theta, spindle, and slow oscillations), LFP was recorded with respect to the GND screw. Electrodes were not moved at least 4 hours before and during the recording day.

Data were collected using a SpikeGadgets data acquisition system (SpikeGadgets LLC). Spike data were sampled at 30 kHz and bandpass filtered between 600 Hz and 6 kHz. LFPs were sampled at 1.5 kHz and bandpass filtered between 0.1 Hz and 400 Hz. The animal’s position and running speed were recorded with an overhead color CCD camera (30 fps) and tracked by color LEDs affixed to the headstage.

### Spike sorting

Single units were sorted using a custom manual clustering program (MatClust, M. P. Karlsson) and were distinguished based on spike width, peak and trough amplitude, and principal components. Cluster quality was measured using isolation distance (*48*), and only well isolated neurons were included for analysis.

Putative CA1 pyramidal cells (43.25 ± 3.06 pyramidal units per epoch) and interneurons were further separated by burst probability, mean firing rate, and trough-to-peak (fig. S6). Burst probability was defined as the proportion of spikes within an epoch that occurred within 6 ms of each other (*49*). All well isolated PFC units were included for analysis (34.36 ± 2.08 units per epoch).

### Sleep state identification

To detect bouts of NREM and REM sleep, animals’ head speed and CA1 LFPs were used as previously described (*19, 50*). During sleep box sessions, times of awake activity and immobility were defined as periods where head speed was greater or less than 4 cm/s, respectively. Candidate NREM sleep bouts were identified as periods where head speed remained <4 cm/s after an immobility period of at least 60 s. NREM and REM sleep were further identified and separated by using a theta to delta ratio metric. Briefly, CA1 LFP (referenced to GND) for each tetrode was bandpass filtered (8-12 Hz for theta; 1-4 Hz for delta) and averaged, and the ratio of theta to delta power was calculated. Periods that exceeded a set threshold (mean + 1 SD) for >10 s were categorized as REM bouts within candidate NREM bouts (fig. S2A).

### LFP event detection

CA1 and PFC ripple detection: Ripple events were detected during immobility periods (<4 cm/s) as previously described (*19*). Tetrodes with recorded cells in CA1 and PFC were used as electrodes for ripple detection. Each LFP from candidate electrodes were bandpass filtered in the ripple band (150-250 Hz), and the envelope power was calculated using a Hilbert transform. Ripple events were detected as contiguous periods where the ripple power exceeded 3 SD of the mean on at least one tetrode (fig. S2B). Events were further refined as periods around the detected event that remained above the mean (start and end times). The amplitude of each ripple event was reported as the number SDs above the mean (*12*). To estimate the frequency of each ripple event, the power in each frequency bin was calculated around each ripple event (±500 ms) using multi-taper time-frequency analysis (Chronux toolbox; http://chronux.org/) and was z-scored relative to a baseline spectrogram. The frequency of each event was defined as the frequency with the highest power in the ripple band (100-250 Hz) between the start and end time of each event (*51*). Furthermore, ripple events in CA1 and PFC were designated as independent or coordinated based on whether cross-region event times (start to end intervals) overlapped. The coordinated events were further subset into leading or lagging events depending on the identity of the leading ripple. Only events where there was a clear temporal relationship were used (i.e. the start and end times of the leading event preceded the start and end times of the lagging event, respectively).

PFC spindle detection: Spindle detection was performed using the same PFC tetrodes used for ripple detection. Similar to ripple detection, LFP from PFC tetrodes were bandpass filtered in the spindle band (12-16 Hz), and the envelope power was calculated using a Hilbert transform. Contiguous periods that exceeded 3 SD of the mean on at least one tetrode were considered spindle events. Events were further refined as periods around the detected event that remained above the mean (start and end times).

PFC slow wave detection: To extract slow waves from PFC LFP, tetrodes that were used to detect PFC ripple events were averaged together and filtered through two independent filters as previously described (*52*). Briefly, the averaged LFP was filtered with a high pass Butterworth filter (0.1 Hz cutoff, 2^nd^ order) followed by a low pass filter (4 Hz cutoff, 5^th^ order). To separate SOs from other, lower amplitude slow waves (i.e. possibly delta oscillations), the following process was employed. Here, we are considering SOs as high amplitude waves in the 0.1-4 Hz range. First all zero crossings were identified and each segment that included an up state between two down states was extracted for further processing. For each epoch, a positive (>85 percentile of peaks) and negative threshold (<40 percentile of troughs) was set for down and up states, respectively. A wave was considered a SO if both the first peak was higher than the positive threshold and the trough was lower than the negative threshold. Slow waves were classified as low amplitude (other) if the trough was lower than the negative threshold but the first peak did not surpass the positive threshold. Furthermore, only events where the time from first peak to trough was 150-500 ms were included. To visualize the separation of SOs and other slow waves, K-Means clustering was performed on the mean peak and trough values from each epoch (fig. S15A).

All of the above events were further subset by periods of NREM sleep for subsequent analysis.

### Ripple rates and ripple rate correlation

Ripple rates were calculated for each animal and epoch in sessions where the total duration of NREM sleep was >30 seconds. This yielded a rate vector for each animal, which was z-scored to account for the differences in ripple rate across animals.

In addition, to examine how changes in either CA1 or PFC ripple rate may drive changes in the occurrence of coordinated events in NREM sleep, the Pearson correlation was taken between the ripple rate vector calculated from all events and the rate vector for coordinated events. Within area (CA1_all_-CA1_coord_ or PFC_all_-PFC_coord_) r values were computed for each animal and compared.

To investigate changes in ripple rate within a sleep session (relative rate change), NREM sleep was split into two halves, 1^st^ and 2^nd^, by dividing the total duration by 2. The ripple rates were calculated for each half, and the relative rate change was calculated as follows:

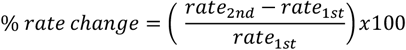

Only epochs with a total NREM duration exceeding 120 seconds (60 s per rate calculation) were included for analysis.

### Event cross-correlation

To examine whether independent and coordinated events are differentially coupled with spindle and SOs, the cross-correlation between event onset time vectors was calculated for each event pair (max time lag 4 seconds, 100 ms time bins) and smoothed with a Gaussian kernel (300 ms). For each comparison, ripple times were used as the reference vector. Thus, positive and negative lag values indicate that the compared LFP event tended to occur before and after ripple events, respectively.

To calculate 95% confidence intervals, both CA1 and PFC ripple times were jittered using the following steps. First, initial jitter values were resampled from the distribution of inter-event intervals separately for CA1 and PFC. Then, these values were randomly assigned negative or positive values to define the direction of the jitter. Finally, a value between 1 and 5 seconds was added to the direction of the jitter before calculating the cross-correlation. This process was repeated 1000 times for each epoch, and the 95^th^ and 5^th^ percentiles were calculated. This time jitter was chosen instead of a circular shuffle since only ripple times during NREM sleep were used for this calculation. A random circular shuffle would have likely shifted ripple times into periods not defined as NREM, yielding inaccurate results.

### Ripple aligned modulation

Single unit ripple modulation in CA1 or PFC was calculated as previously described (*17*). This analysis was restricted to neurons that fired >50 spikes cumulatively across all peri-ripple windows (±2 s). For each candidate neuron, an averaged ripple-triggered (CA1 or PFC ripple) time histogram (ripple-PSTH) was calculated. For comparison, 1000 shuffled ripple-PSTHs were generated by circularly shifting the spikes around each ripple event by a random amount. The determine whether a neuron was significantly modulated, the squared difference between the real ripple-PSTH and the shuffled data in the 0-200 ms window after ripple onset was calculated. In the case of CA1 modulation during independent PFC ripples, a larger window spanning ±200 ms around ripple onsets was used due to the observed CA1 population response aligned to PFC ripples (fig. S14B). If this real value exceeded 95% of the shuffled values, the neuron was categorized as significantly modulated. Furthermore, excited (EXC) and inhibited (INH) neurons were separated based on whether the modulation in ripple response window (either -200 to 200 or 0 to 200 ms) was positive (EXC) or negative (INH). Modulation index (MI) for each cell was calculated as the mean response in the background window (−600 to -200 ms prior to event onset) subtracted from the mean response in the response window.

### Spectral analysis

Wavelet spectral analyses were utilized to calculate the ripple aligned power spectra in PFC (2– 350 Hz, Morlet wavelets). For each epoch, the PFC tetrode with the greatest number of detected PFC ripples was used. The power at each level of the wavelet transform was calculated around each ripple event (±500 ms) and was z-scored relative to a baseline spectrogram calculated for the entire epoch. For each animal, spectrograms from all sleep epochs were averaged (fig. S3A).

### Ripple triggered spindle power

To determine whether there was a difference in spindle power during independent vs coordinated events, the envelope magnitude for each PFC spindle detection tetrode was z-scored and the maximum envelope power during each ripple event was calculated and averaged across electrodes.

### Theta phase-locking

Theta phase-locking during active behavior on the W-Track was quantified using previously developed methods (*53*). The CA1 reference tetrode located in corpus callosum was filtered in the 6-12 Hz range and was used to extract phase information, since theta phase has been reported to be consistent at different depths above the CA1 pyramidal layer (*54*). Spikes were restricted to theta periods where the animals’ speed was >5 cm/s, and spikes were assigned a phase between 0-360°. Only neurons with at least 100 spikes during theta periods were included in this analysis. The Rayleigh test was used to determine whether the phase distribution of each cell deviated from circular uniformity (p < 0.05 criterion). Subsequently, for each theta modulated cell, phase locking strength was measured using two separate methods, the circular concentration parameter, kappa (κ), and pairwise phase consistency (PPC) (*55*). κ was estimated by fitting the phase distributions to a von Mises distribution, and PPC was calculated by taking the dot product of the angular difference between each pair of phase values then averaging. To test for phase preference differences, the distributions were compared using a nonparametric Watson’s U^2^ test.

### Single unit properties

Linearized spatial maps during active behavior on the W-Track were calculated during periods of high mobility (>5 cm/s; all SWR times excluded) at positional bins where occupancy was >20 ms (*17*). Briefly, each rat’s 2D position was projected onto a linearized skeleton of the W-Track and was classified as belonging to one of the four possible trajectories. Then, the linearized spatial maps were calculated by dividing the binned spike count by the occupancy in each 2 cm positional bin and smoothed with a Gaussian kernel (4 cm).

For each epoch, mean theta firing rate was calculated by dividing the number of spikes that occurred during mobility periods (>5 cm/s) by the total duration of mobility. Similarly, NREM firing rate was calculated by dividing the number of spikes during NREM (excluding SWR times) by the total duration.

To calculate the number of place fields for each CA1 neuron, 5,000 null linearized firing rate vectors were generated by randomly sampling each cell’s spikes using the smoothed occupancy as the probability density function. The resulting spike vector (concatenated across all four trajectories) was smoothed with a Gaussian kernel (4 cm) and divided by the smoothed occupancy. Each bin that exceeded the 99^th^ percentile of the null distribution was determined as having selective firing within that bin. To be considered a place field, at least 5 consecutive bins (10 cm) exceeding the 99^th^ percentile was required (*32*). Field width was calculated as the mean width across all place fields for each cell.

Spatial information, I (bits/spike), was calculated using the following formula:

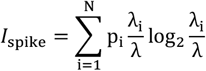

where i = 1, …, N represents the spatial bin, *p* is the occupancy probability of bin *i*, and λ is the mean firing rate of the neuron.

To calculate the field stability of modulated CA1 neurons, the Pearson correlation of the concatenated linearized firing fields was taken between the two W-Track epochs flanking the sleep epoch during which the cells were modulated.

### Single unit SWR activity

The firing rate gain of each neuron was calculated by dividing the firing rate during CA1 SWR events by the mean rate across all NREM bouts in an epoch. Additionally, to exclude the contribution of SWR firing to the baseline NREM firing rate, the mean rate was calculated by excluding spiking activity across all SWR events during NREM sleep. To determine SWR burst probability for each cell, the number of SWRs where burst spikes were detected (at least 2 spikes <6 ms apart) was divided by the total number of SWR events in which that cell was active. Similarly, SWR spike latency was estimated by taking the average latency to first spike during each SWR event the cell was active. For quartile analysis, cells were sorted by modulation index and split into 4 equal groups. The corresponding SWR spiking metrics were then averaged and plotted for each group. To determine whether there was a relationship between the modulation indices and burst probability or spike latency, the Pearson correlation coefficient was calculated.

### Event aligned multi-unit activity

To characterize population activity aligned to LFP events, spike trains of each neuron were binned into 5 ms time windows. Activity was aligned to PFC spindles, SOs, ripples, and CA1 ripples. The event aligned multi-unit activity was then normalized by the mean population firing rate during NREM sleep.

### Ripple coactivity

To quantify ripple coactivity during independent and coordinated events, a pairwise coactivity z-score was calculated as previously described (*56*). Briefly, this coactivity measure estimates how likely two cells will fire together during ripple events and is normalized by the incidence of spiking independent of the other cell:

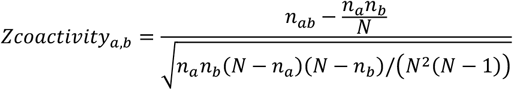

where *N* is the number of ripple events, *n*_*a*_ and *n*_*b*_ are the number of events where cell *a* and cell *b* were active, respectively, and *n*_*ab*_ is the number of events where both cells were active. Epochs where there were >20 independent and coordinated events were included in the analysis.

### Assembly detection

To detect cell assemblies in CA1 and PFC populations, a previously described method was used (*57, 58*). Significant co-firing patterns in CA1 or PFC were detected using a method based on principal and independent component analyses. For each W-Track epoch, the spike trains of each neuron were binned into 20 ms time windows (excluding CA1 ripple times) and Z-scored to eliminate any biases due to differences in firing rates. Principal component analysis was then applied to the Z-scored spike matrix (Z). The resulting eigenvalue decomposition of the correlation matrix (C) was given by:

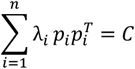

Where *p*_*i*_ is the *i*^*th*^ principal component of C and *λ*_*i*_ is the corresponding eigenvalue. Note that 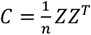 is the correlation matrix of Z. To determine the number of significant coactivity patterns in the data, the Marcenko-Pastur law was used to calculate a threshold eigenvalue λ_*max*_, which was given by 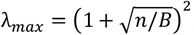, where *n* is the number of recorded units and *B* is the total number of bins. Note that an eigenvalue exceeding *λ*_*max*_ indicates that the corresponding cell assembly explains more correlation than expected if the activity of neurons was independent of each other. The number of eigenvalues above λ_*max*_, *N*_*A*_, represents the number of significant patterns found in the data. These principal components were then projected back onto the original z-scored spike data:

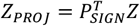

where *P*_*SIGN*_ is the *nXN*_*A*_ matrix with *N*_*A*_ columns. Independent component analysis (ICA, RobustICA) (*59*) was then used to identify patterns that were maximally independent from each other. ICA was applied to the matrix *Z*_*PROJ*_ to find an *N*_*A*_*XN*_*A*_ unmixing matrix, *W*, which was used to obtain the weights of each cell in each assembly.

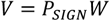

Since the sign of the output of ICA is arbitrary, the signs of the weight vector were set such that the highest absolute weight was set to positive. Cells with weights greater than the mean + 2SD were considered member cells. Furthermore, field similarity for member vs non-member cell pairs was calculated by taking the Pearson’s correlation between the concatenated linearized firing fields (fig. S9A).

In addition to the assembly weights, a contribution metric, *I*, was calculated to assess the relative influence of EXC and INH cells on assembly reactivation during sleep. To quantify the contribution of a cell, *c*, the reactivation strength of each assembly, *k*, was recalculated after omitting the spike train of that cell, 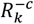, to yield two separate reactivation strength vectors, 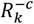 and *R*_*k*_. The contribution was then calculated by:

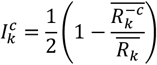

where ½ is a normalization factor such that the sum of the contributions equals 1 (*57*).

### Assembly reactivation strength

To assess the reactivation of these detected assemblies during sleep sessions, the expression strength of each pattern over time was given by:

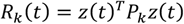

*R*_*k*_(*t*) is the reactivation strength of assembly *k* at time *t, z*(*t*) is the z-scored firing rate vector for all neurons at time *t*, and *P*_*k*_ is the projection matrix for assembly *k*, constructed by taking the outer product of its weight vector *v*_*k*_. Additionally, the diagonal of the projection matrix was set to zero to reduce the influence of highly active neurons. Thus, only patterns of coactivity are detected as periods of high reactivation strength. For CA1-PFC assembly analysis, within region correlations in the projection matrices were also set to 0 to measure only the coactivity between areas. Also, only CA1-PFC assemblies that had at least one member cell in each area were included for analysis. Only sleep sessions with a preceding run session to use as a template were included for reactivation analysis (sleep sessions 2-9).

### Assembly coactivation rate

To investigate CA1-PFC assembly coactivity during independent and coordinated CA1 ripple events, a coactivation rate was calculated by first defining discreet periods of high reactivation (*R* > 5). For each ripple event, the number of CA1 and PFC assemblies that were active was calculated separately for each area. The events were then subset to include events where both CA1 and PFC assemblies were active. The minimum number of active assemblies across areas was used as the number of coactivation events (i.e., CA1-2, PFC-3 would be 2 coactivation events). The total number of events across all ripples was divided by the cumulative ripple time to obtain a coactivation rate per epoch.

### Ripple triggered assembly strength

The reactivation strength of each assembly aligned to ripple onsets was determined by calculating the mean, Z-scored reactivation strength in a ±8 s window around the ripple triggered events. Then, the average across the peak bins (±200 ms) was calculated to obtain a single reactivation strength value for each assembly. Using this method to calculate overall reactivation strength yielded a gradient of strengths as expected (fig. S10). For epoch grouped analysis, epochs 2-4, 5-6, and 7-9 were used for early, middle, and late sleep sessions, respectively.

### Assembly reinstatement

To assess assembly reinstatement from one W-track session to another, the above assembly reactivation analysis was performed, but instead, cell identity was matched across three consecutive epochs (run-sleep-run) to measure the same assembly over time. The epochs were matched such that each W-track session acted as the template epoch, except for the 8^th^ W-track session. Reinstatement was defined as the difference in the average expression strength during the two compared exposures (2^nd^ exposure minus 1^st^ exposure) (*58*).

### Replay analysis

Bayesian decoding for replay analysis was performed as previously described (*60*). Only sleep sessions with a preceding run session to use as a template were included for replay analysis (sleep sessions 2-9). A memoryless (uniform prior probability) Bayesian decoder was built for each of the four trajectory types to estimate the probability of animals’ position given the observed spikes (Bayesian reconstruction; or posterior probability matrix):

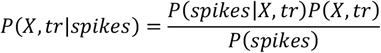

where *X* is the set of all linear positions on the track for different trajectory types (*tr* ∈ {center-to-right, right-to-center, center-to-left, left-to-center}), and we assumed a uniform prior probability over *X* and *tr*. Assuming that all *N* place cells active during a candidate event fired independently and followed a Poisson process:

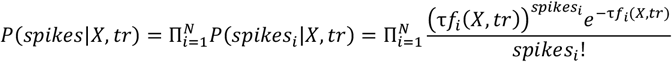

where *τ* is the duration of the time window (15 ms), *f*_*i*_(*X,tr*) is the expected firing rate of the *i*-th cell as a function of sampled location *X* and trial type *tr*, and *spikes*_*i*_ is the number of spikes of the *i*-th cell in a given time window. Therefore, the posterior probability matrix can be derived as follows:

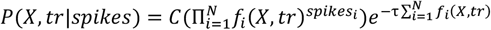

where *C* is a normalization constant.

We identified replay during SWRs based on the Bayesian decoder described above. First, candidate events were defined as the SWR events longer than 50 ms during which ≥ 5 place cells fired. Each candidate event was then divided into 15-ms non-overlapping bins, and the posterior probability matrix was computed based on the Bayes’ rule. To determine replay significance, we employed the two following shuffle-based methods.

### Replay significance (Monte Carlo method)

In this first method, the *p*-value was calculated based on the Monte Carlo shuffle (*61*). First, we drew 10,000 random samples from the posterior probability matrix for each decoded bin and assigned the sampled locations to that bin. Linear regression was then performed on the bin number versus the location points. The resulting *R*-squared was compared with 1,500 regressions, in which the time bins were shuffled). A candidate event with p < 0.05 based on the time shuffle was considered as a replay event. Since the shuffling procedure measured how well the decoded positions along SWR time matched a behavioral trajectory, we considered the trajectory with the lowest *p*-value from the shuffling procedure as the replay trajectory, and its *R*-squared was reported as replay quality (fig. S12).

### Replay Significance (Weighted Correlation vs. shuffle)

Additionally, a sequence score was computed for each candidate replay event as the correlation between position and time, weighted by the posterior probability (*32, 62*). A weighted mean was calculated for each event by:

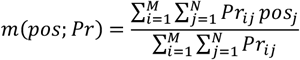

The weighted covariance by:

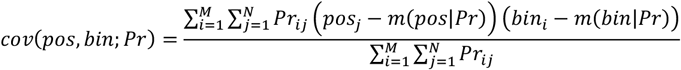

Where *pos*_*j*_ is the *jth* spatial bin, *bin*_*i*_ is the *ith* time bin, *M* is the number of time bins, *N* is the number of spatial bins, and *Pr*_*ij*_ is the posterior probability for the spatial bin in that time bin.

Finally, the weighted correlation was given by:

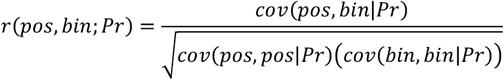

Subsequently, 1,000 shuffled spatial template matrices were constructed by circularly shifting each cell’s template independently. The entire Bayesian decoding process was performed using these shuffled templates to yield a null distribution of weighted correlation values for each candidate replay event. An event with *p* < 0.05 based on the spatial template shuffle was considered as a significant replay event. Additionally, for each event, a sequence score (*rZ*) was calculated by:

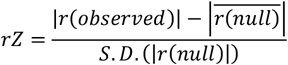

Where S.D. is the standard deviation of the null distribution. Note that the *rZ* value does not signify directionality of the replay event.

Jump distance for each significant event was reported as the distance between the maximum decoded probability of two adjacent time bins. Each pair of time bins yielded a distance value unless there were bins with no spiking activity. The maximum of these values was taken and normalized by dividing by the length of the corresponding trajectory.

### Per cell contribution (PCC) analysis

PCC was calculated for each cell to assess the influence of a cell’s participation in a replay event (*32*). For each candidate replay event, each cell’s place representation was circularly shuffled separately such that only one cell’s spatial representation was shuffled at a time. Each cell’s representation was shuffled 1000 times to calculate an *rZ*_*cellshuffle*_ value as above. Then, PCC was given by:

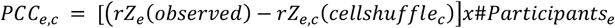

Where ‘#*Participants*_*e*_’ is the number of cells active during event *e*. The average PCC was calculated for cells that were active in >10 significant replay events per epoch. For the calculation of PCC difference between coordinated and independent events, the cell was required to be active in >5 events of each type per epoch.

### Rank-order correlation

To evaluate the consistency with which populations of neurons in CA1 fired in sequence, we calculated the rank-order correlation between each event and its template as previously described (*23, 63*). For each SWR event with >5 cells active, the spiking activity (both pyramidal cells and interneurons included) was reduced to each unit’s first spike time, chronologically ordered, and normalized from 0 to 1. Then, a cross-validated template-based method was used to calculate sequence similarity between each event and its template. For each event, a template was created by averaging the normalized rank of each cell across all events excluding the event in question. The rank correlation was reported as the Pearson correlation coefficient between the event and template. Templates were constructed separately for independent and coordinated events, utilizing only events of one type.

### Population dimensionality during SWRs

To investigate the dimensionality of population activity during independent and coordinated SWR events, principal component analysis (PCA) was performed on separate spiking activity matrices. Sessions with >20 units (pyramidal and interneurons) were included for analysis. Since independent events outnumbered coordinated events, we only included sessions where there were >100 events of each type. Then, we randomly resampled each activity matrix 100 times (50 samples at a time) and applied PCA to each matrix to obtain the variance explained by each component (n components = n cells). We considered the population dimensionality to be the proportion of components required to explain >90% of the variance.

### Ripple type prediction

Ripple type (independent or coordinated) in CA1 and PFC was predicted by fitting a linear model (MATLAB, fitclinear) to the areas’ spike count data. Since the independent events largely outnumbered the coordinated events, the event counts were equalized prior to training by randomly excluding a subset of independent events. The binary decoders were built using 10-fold cross validation, and prediction accuracy was assessed by comparing performance against models constructed with randomly permuted spike matrices (n = 1000). To account for the dropped events, this entire process was repeated 10 times for each animal and epoch. This method was also used to predict different coordinated events (CA1 or PFC leads), but instead, both CA1 and PFC spiking activity was used.

### Replay probability during SOs

To calculate the probability of CA1 replay events during the down or up state of SOs, the occurrence of replay start times of each significant event (independent, coordinated, or all) was determined and divided by the total number of replay events in NREM sleep for each epoch.

### Peri-SO event probability

To investigate the sequential occurrence of LFP events during NREM sleep, the fold change occurrence around SO troughs was calculated for PFC spindles, ripples, and CA1 ripples. Event start times were separately binned in 20 ms time bins in a response window ±2 seconds (resp) centered around SO troughs (up states). The event probability in each bin was then calculated, smoothed with a 60 ms Gaussian kernel, and compared to the mean probability across all bins in the background window -3 to -2 seconds (bck) around SO trough. The fold change in each bin was then calculated by:

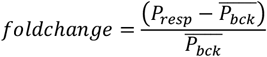

Since this approach results in a single fold change vector, we utilized a resampling method to determine whether the resultant peaks were not due to chance. The occurrence matrix was randomly sampled (100 samples at a time), and the probability fold change was calculated as above with the corresponding background bins. This process was repeated 50 times to obtain a distribution of values in each time bin. Then, the bin in which the peak fold change occurred was compared to 0 with a ttest.

To compare the timing of events aligned to SO troughs, the distributions of all event times within ±250 ms of SO troughs were compared. To further validate this timing result, distributions of the times of peak fold change occurrences around SO troughs were compared by calculating event fold change separately for each epoch and animal.

To investigate the SO phase preference of spindles and ripples, phase information for each discreet SO event (upstate and downstate) was extracted. Each LFP event (event onset) that occurred within the start/end interval of SOs was assigned a phase. Then, a circular von Mises probability density function was computed based on parameters (preferred direction and concentration parameter) estimated from the distribution of phase angles. To test for phase preference differences for spindles and ripples, the distributions were compared with Watson’s U2 test.

### Peri-SO CA1 and PFC reactivation strength

To investigate the temporal dynamics of CA1 and PFC reactivation during these events, we looked at CA1 lagging events surrounding SO troughs. Reactivation strength of each assembly was aligned to SO troughs where there was a CA1 lagging ripple event within ±250 ms of a SO trough. As a comparison, each assembly was circularly shuffled, and the peak reactivation bin of the real data was compared to the same bin of the shuffled data. In addition, to compare the relative reactivation times of CA1 and PFC assemblies, the peak of the mean reactivation strength for each was taken in the 0-200 ms time window surrounding SO troughs, and the timing distributions were compared. The same process was performed for CA1 leading events, but in this case, timing differences were assessed using a ±100 ms window around SO troughs. The windows were determined by examining the reactivation profiles surrounding SO troughs.

To compare CA1 assembly suppression during independent PFC ripples to CA1 reactivation during SO associated coordinated events where CA1 ripples lag, the suppression strength of each assembly in a ±200 ms window around PFC ripples was calculated as in “*Ripple triggered assembly strength*.” Then, the mean reactivation strength in a 0-200 ms window around SO troughs was compared to the suppression during PFC ripples for each assembly.

### Histological verification of recording sites

At the completion of each experiment, electrolytic lesions were made by sending current (30uA) down 3 electrodes per tetrode for 3-4 seconds each. 24 hours later, animals were sacrificed by intraperitoneal injection of Euthasol and subsequent intracardial perfusion with 0.9% saline followed by 4% formaldehyde. The skull, with the attached drive, was submerged in 4% formaldehyde for 24 hours before tetrode retraction and brain extraction. Brains were stored in a 4% formaldehyde and 30% sucrose solution for at least 1 week before slicing with a freezing microtome (Leica) at 50uM. Slices were Nissl stained and imaged at 4x on a brightfield microscope (Keyence) and stitched together to generate composite images for lesion localization.

### Statistical Analyses

All statistical analyses were performed with MATLAB (MathWorks, Natick, MA; R2022a) functions or custom scripts. No specific analysis was performed to predetermine sample size, but the number used in this study is similar to or larger than typically used. Nonparametric, two-tailed statistical tests (Wilcoxon rank-sum and signed-rank, for unpaired and paired data, respectively) were used throughout the paper unless noted. P < 0.05 was considered the cutoff for statistical significance. Boxplots show the median (center line), interquartile range (box), and the furthest points not considered outliers (whiskers). Although not displayed in some cases, outliers were included in all statistical analyses. Unless otherwise noted, values and errors in the text denote means ± SEM. p-values in the text are reported as follows: p > 0.05, *p < 0.05, **p < 0.01, ***p < 0.001.

**Fig. S1.**
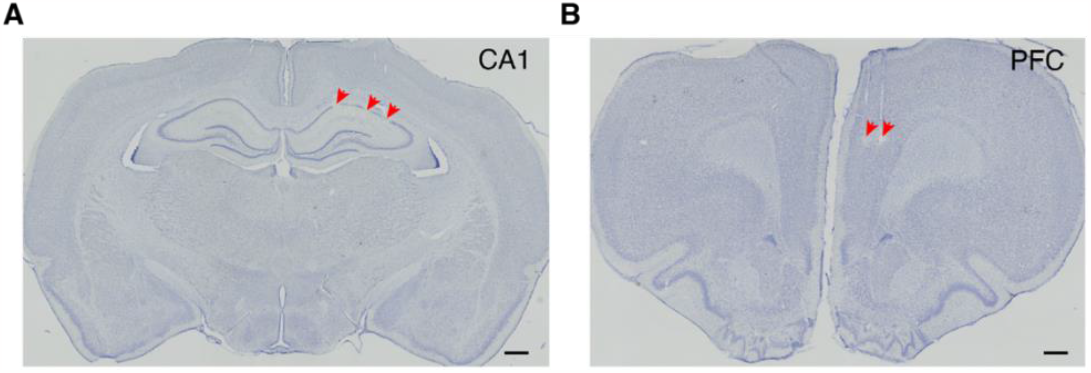
Histological verification of recording locations. Lesions were visualized with a Nissl stain. Red arrows show example lesion sites in (**A**) dorsal CA1 and (**B**) the prelimbic region of PFC. Scale bars, 500 μm.

**Fig. S2.**
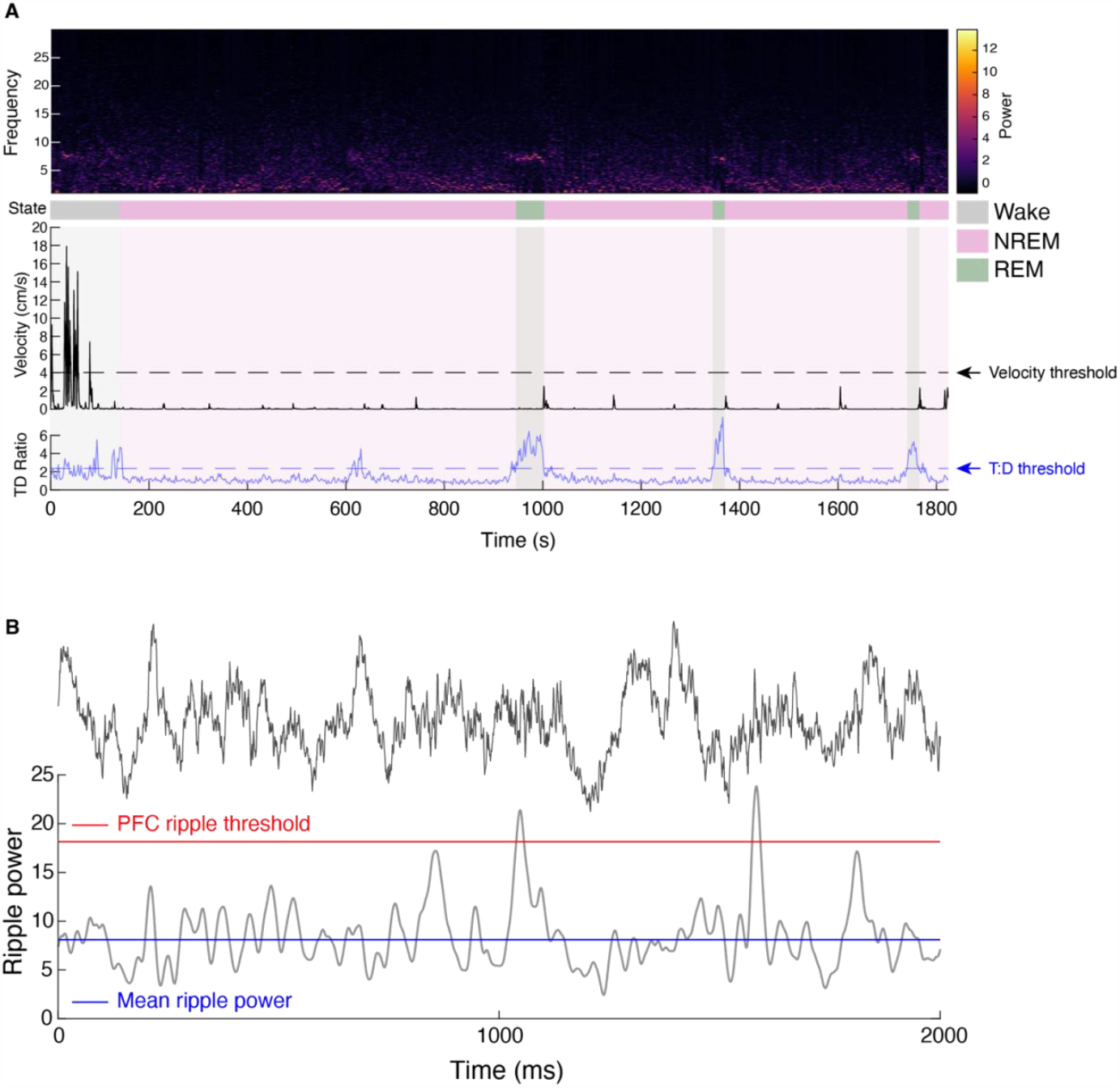
Sleep state designation and detection of PFC ripple events. (**A**) Example sleep session illustrating the method used to delineate behavioral state using hippocampal LFP and velocity. Note the increase in theta band activity and the absence of movement during periods of REM sleep (TD ratio: theta to delta ratio). (**B**) Method used to detect ripples in PFC. For each tetrode, ripples were extracted using a threshold of 3SD above the mean ripple power. Similar procedures were used to extract spindle events.

**Fig. S3.**
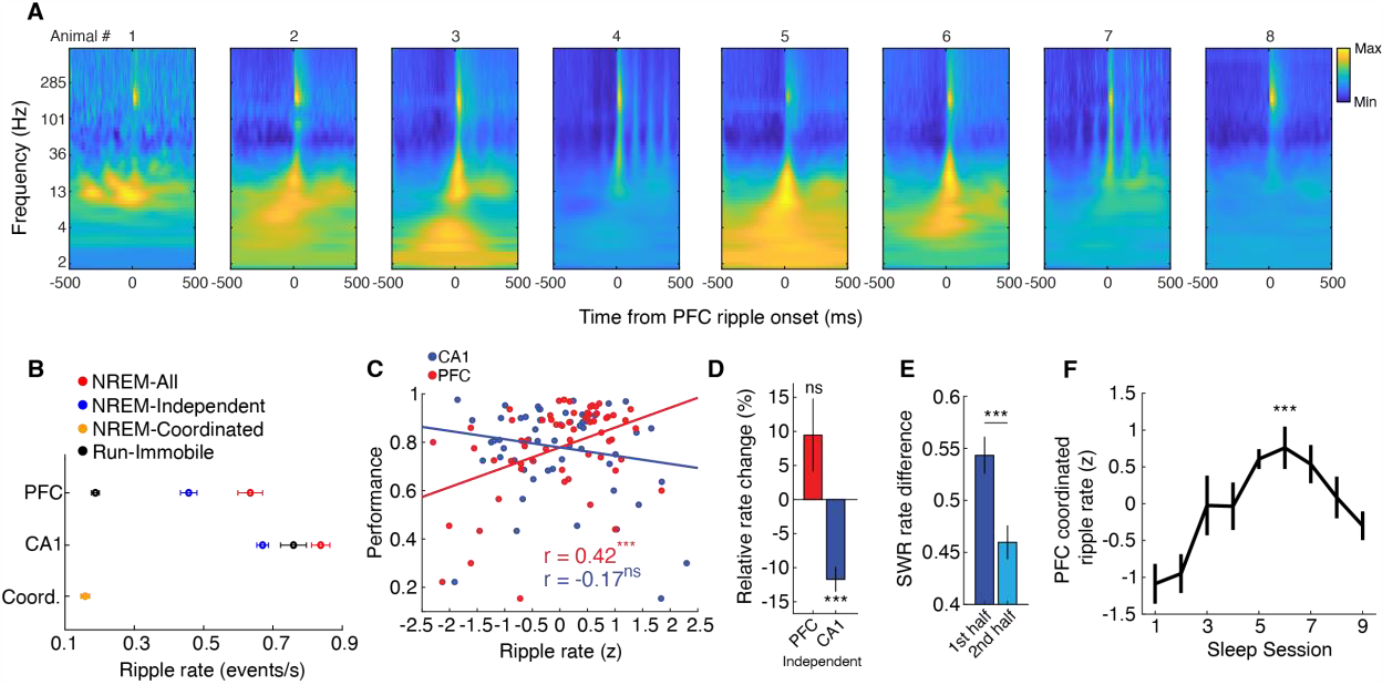
PFC ripple triggered spectrograms and ripple rates. (**A**) PFC ripple triggered spectrograms for each animal. Note the prominent spindle band activity (12 – 16 Hz) during these events in most of the animals. (**B**) Ripple rates in PFC and CA1 for the different events (CA1 all = 0.84 ± 0.03; CA1 independent = 0.67 ± 0.02 Hz; CA1 run = 0.76 ± 0.04 Hz; PFC all = 0.63 ± 0.04 Hz; PFC independent = 0.46 ± 0.02 Hz; PFC run = 0.19 ± 0.01 Hz; Coordinated = 0.16 ± 0.01 Hz). (**C**) Correlation between independent ripple rate and overall performance in the run session following the sleep session in which the rate was calculated (left; PFC, r = 0.42, ***p = 8.30×10^−4^; CA1, r = -0.17, p = 0.18). The right panel shows the difference between the correlation coefficients for PFC vs CA1 (PFC, r = 0.42 ± 0.12; CA1, r = -0.17 ± 0.13, ***p = 8.68×10^−4^, z-test for comparison of correlation coefficients). (**D**) Change in ripple rate relative to the first half of NREM sleep. Independent SWR rate decreased in the second half of NREM sleep during each epoch, whereas PFC ripple rate remained consistent (PFC = 9.46 ± 5.35, p = 0.08, CA1 = -11.72 ± 1.82, ***p = 1.71×10^−8^, ttest vs 0). (**E**) SWR rate difference between independent and coordinated events during the first and second half of NREM sleep. The drop in independent SWR rate shown in (**D**) shifted the balance of independent and coordinated events (First half = 0.54 ± 0.02, Second half = 0.46 ± 0.02, ***p = 2.62×10^−6^, Wilcoxon signed-rank). (**F**) Z-scored coordinated PFC ripple rate over epochs (F = 5.15, ***p = 3.47×10^−4^, Repeated measures ANOVA, main effect of time). Only PFC rate is shown here since the curves for PFC and CA1 largely overlap due to the requirement of being coordinated. There were slight differences since multiple ripples in one area could be coupled with a single ripple in the other.

**Fig. S4.**
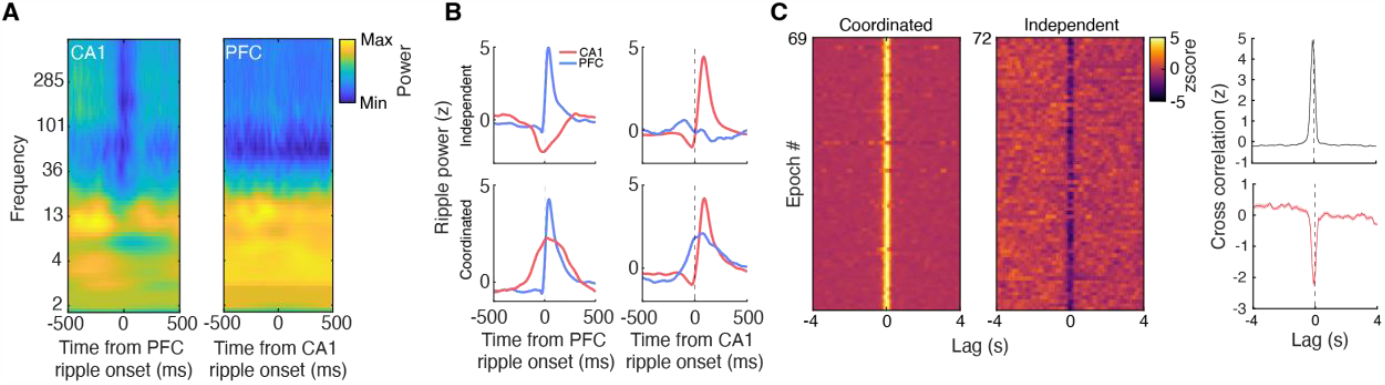
Validation of independent and coordinated events in CA1 and PFC. (**A**) Independent ripple triggered spectrograms showing the absence of ripple band power in the opposing area. (Left) Ripple power in CA1 aligned to independent PFC ripples. Note that independent PFC ripples tend to be associated with attenuated power in multiple frequency bands in CA1. (Right) Ripple power in PFC aligned to independent CA1 ripples. (**B**) Z-scored ripple power in CA1 and PFC during independent and coordinated events. Note the decrease in ripple power in CA1 during independent PFC ripples. (**C**) Cross-correlation between independent and coordinated ripples in CA1 and PFC. Each row represents one epoch from an animal. The number of epochs differ since there were sleep sessions with no coordinated events. The maximum number of epochs is 72 (8 animals x 9 epochs).

**Fig. S5.**
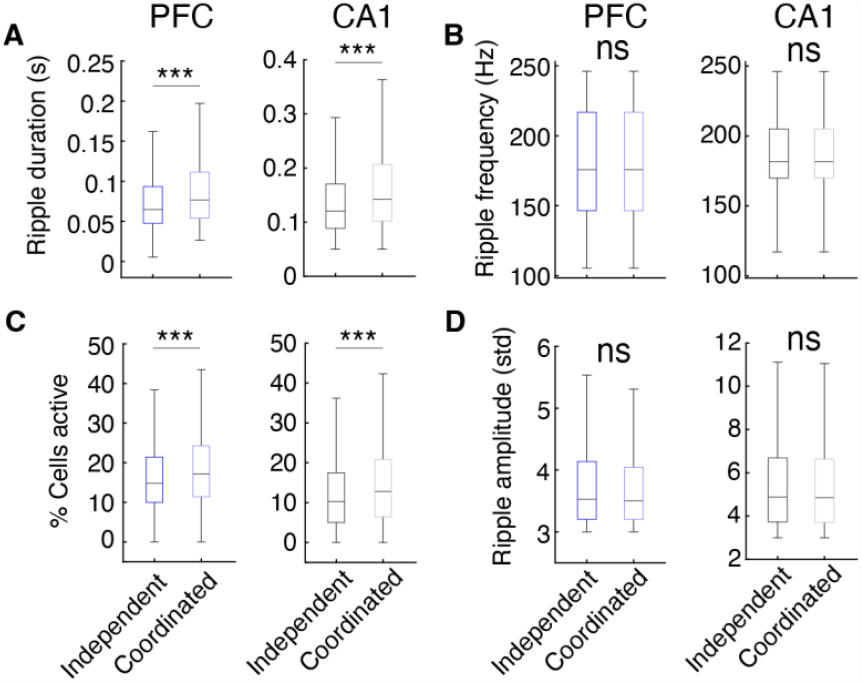
Differences between independent and coordinated ripple events. (**A**) Ripple duration (PFC: Independent = 0.08 ± 2.58×10^−4^ s; Coordinated = 0.10 ± 5.32×10^−4^ s, ***p = 1.06×10^−40^; CA1: Independent = 0.14 ± 4.20×10^−4^ s; Coordinated = 0.17 ± 1.2×10^−3^ s, ***p = 4.26×10^−167^). (**B**) Ripple frequency (PFC: Independent = 183.19 ± 0.17 Hz; Coordinated = 183.80 ± 0.27 Hz, p = 0.16; CA1: Independent = 179.82 ± 0.22 Hz; Coordinated = 180.13 ± 0.42 Hz, p = 0.52). (**C**) Percentage of cells active during ripple events (PFC: Independent = 16.22 ± 0.06%; Coordinated = 18.64 ± 0.09%, ***p = 8.56×10^−99^; CA1: Independent = 12.83 ± 0.05%; Coordinated = 15.22 ± 0.11%, ***p = 1.56×10^−82^). (**D**) Ripple amplitude (PFC: Independent = 3.82 ± 0.01 SD; Coordinated = 3.77 ± 0.01 SD, p = 0.08; CA1: Independent = 5.37 ± 0.01 SD; Coordinated = 5.29 ± 0.02 SD, p = 0.34). Wilcoxon rank sum tests.

**Fig. S6.**
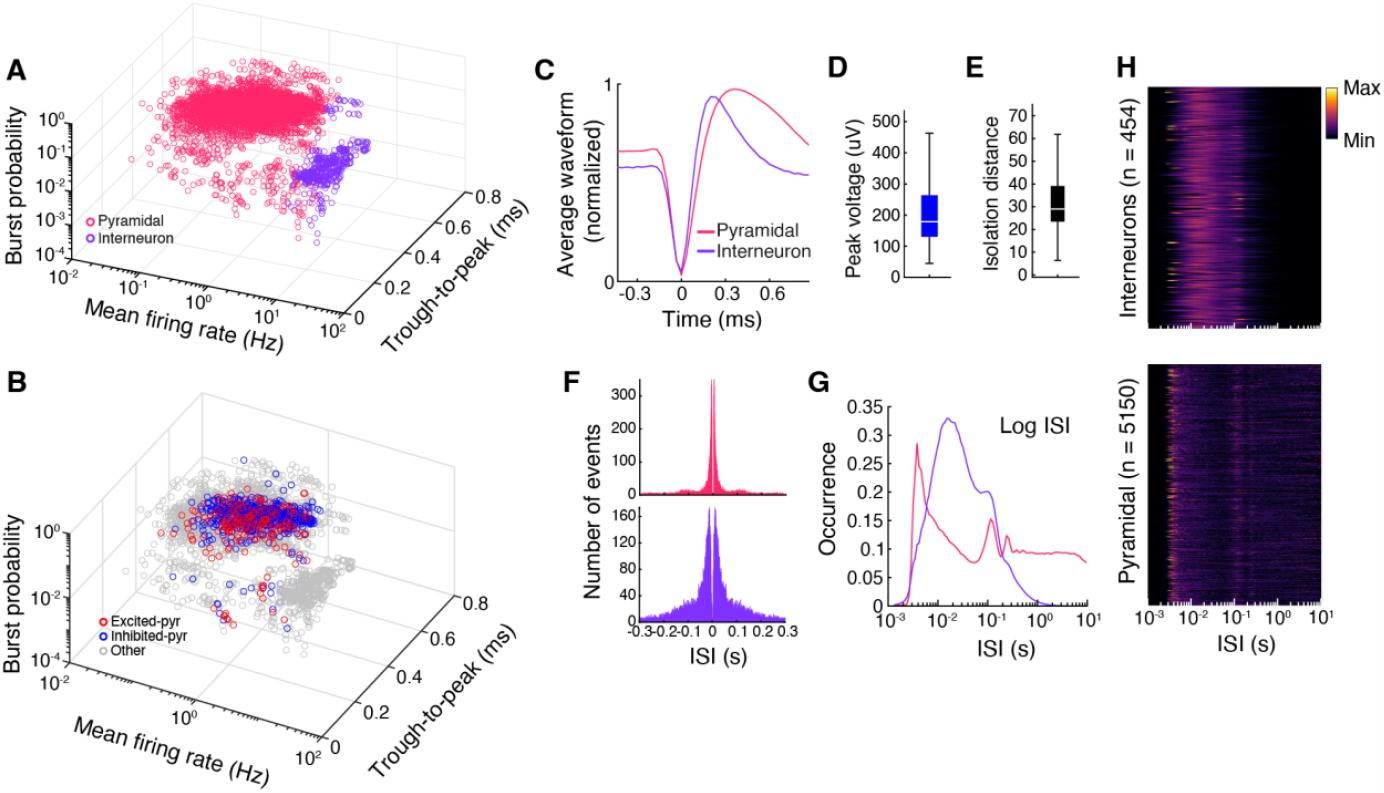
Classification of CA1 pyramidal cells and interneurons. (**A**) To separate populations in CA1 into putative pyramidal cells and interneurons, burst probability, mean firing rate, and trough-to-peak were used. (**B**) Same plot as in (**A**) with CA1 EXC and INH pyramidal cells highlighted to illustrate the overlap within the pyramidal cell cluster. (**C**) Average normalized waveforms for pyramidal cells and interneurons. (**D**) Average peak voltage for each cell measured on the electrode with the highest amplitude (mean = 210.68 ± 1.70 uV). (**E**) Isolation distance of each cell cluster determined by the Mahalanobis distance to the nearest cluster (34.91 ± 1.07). (**F**) Example inter-spike interval (ISI) distributions for an example CA1 pyramidal cell and interneuron. (**G**) Log ISI distributions for pyramidal cells and interneurons. (**H**) ISI heat maps for all evaluated neurons (n = WTrack run + NREM sleep).

**Fig. S7.**
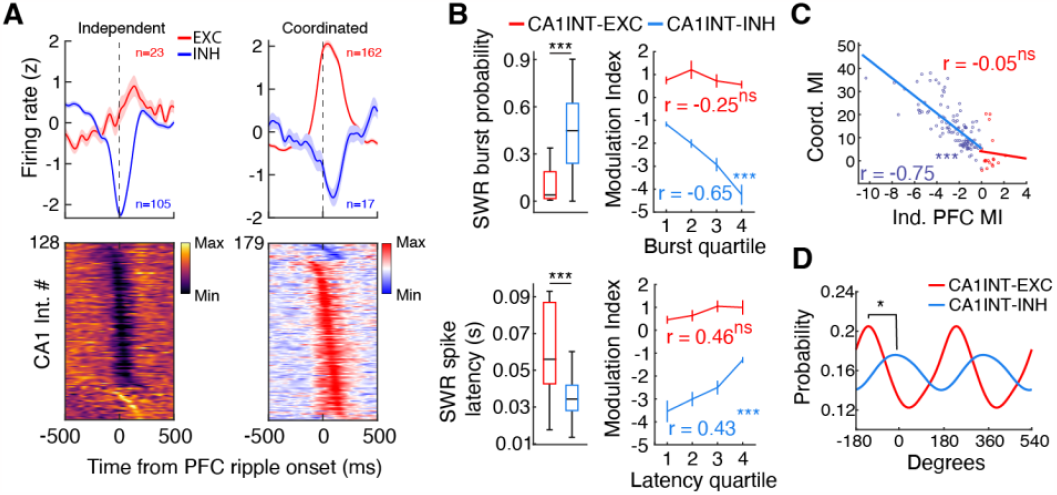
Characterization of the modulation state of putative CA1 interneurons. (**A**) Similar to putative pyramidal cells, interneurons in CA1 were predominantly suppressed in response to independent PFC ripple events. (**B**) Burst probability and spike latency during coordinated SWRs for INH and EXC interneurons (Burst probability: INH = 0.45 ± 0.02; EXC = 0.19 ± 0.08, ***p = 9.43×10^−5^; Spike latency: INH = 0.036 ± 0.001 s; EXC = 0.061 ± 0.007, ***p = 4.37×10^−4^, Wilcoxon rank sum). Similar to CA1 INH pyramidal cells, the modulation state of INH interneurons was predictive of intra-SWR firing dynamics (Burst probability: INH, r = -0.65, ***p = 5.22×10^−14^; EXC, r = -0.25, p = 0.34; Spike latency: INH, r = 0.43, ***p = 5.15×10^−6^; EXC, r = 0.46, p = 0.06). (**C**) Relationship between modulation during coordinated CA1 ripples and independent PFC ripples (INH: r = -0.75, ***p = 5.91×10^−20^; EXC: r = -0.05, p = 0.84). (**D**) Preferred hippocampal theta phase for INH and EXC interneurons (U^2^ = 0.21, *p = 0.03, Watson’s U^2^ test).

**Fig. S8.**
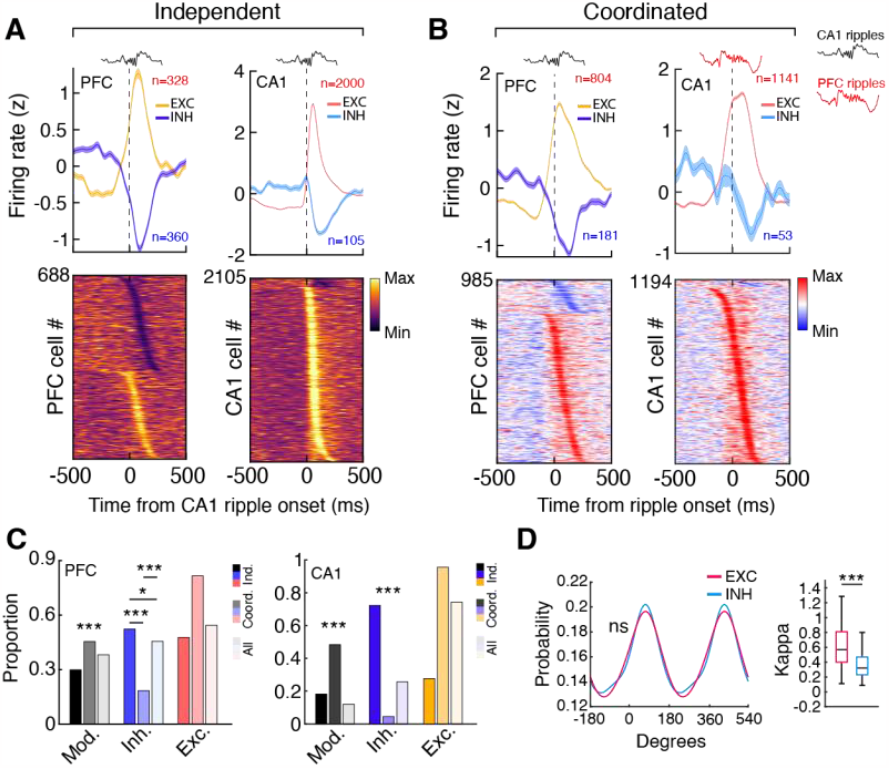
Modulation state of CA1 and PFC is dependent on the separation of ripple events. (**A**) Modulation of PFC and CA1 neurons aligned to independent CA1 SWRs. (**B**) Modulation of PFC and CA1 neurons during coordinated ripples, aligned to ripple onset in the opposing area. (**C**) Proportion of overall modulated cells, INH cells, and EXC cells in PFC (left) and CA1 (right) in response to ripple events in the opposing area. The proportions (PFC: Independent = 0.30, Coordinated = 0.45, All = 0.38; CA1: Independent = 0.18, Coordinated = 0.48, All = 0.12) for overall modulation differed across all three conditions for both PFC and CA1 (PFC: Independent vs Coordinated, ***χ***^2^ = 116.0, ***p = 9.50×10^−27^; Independent vs All, ***χ***^2^ = 34.4, ***p = 8.89x10^−9^; Coordinated vs All, ***χ***^2^ = 24.6, ***p = 1.42×10^−6^; CA1: Independent vs Coordinated, ***χ***^2^ = 544.66, ***p = 3.65×10^−120^; Independent vs All, ***χ***^2^ = 36.6, ***p = 2.92×10^−9^; Coordinated vs All, ***χ***^2^ = 752.79, ***p = 1.99×10^−165^). The relative proportions of INH and EXC cells (PFC INH: Independent = 0.52, Coordinated = 0.18, All = 0.46; PFC EXC: Independent = 0.48, Coordinated = 0.82, All = 0.54; CA1 INH: Independent = 0.72, Coordinated = 0.04, All = 0.26; CA1 EXC: Independent = 0.28, Coordinated = 0.96, All = 0.74) also differed depending on the ripples analyzed (PFC INH: Independent vs Coordinated, ***χ***^2^ = 213.4, ***p = 5.03×10^−48^; Independent vs All, ***χ***^2^ = 6.99, *p = 0.02; Coordinated vs All, ***χ***^2^ = 158.2, ***p = 5.62×10^−36^; CA1 INH: Independent vs Coordinated, ***χ***^2^ = 880.47, ***p = 3.46×10^−193^; Independent vs All, ***χ***^2^ = 161.04, ***p = 1.34×10^−36^; Coordinated vs All, ***χ***^2^ = 133.80, ***p = 1.21×10^−30^). p-values were Bonferroni corrected for multiple comparison. (**D**) Preferred hippocampal theta phase for INH and EXC CA1 pyramidal cells (Watson’s U^2^ test, U^2^ = 0.1045, p = 0.25) and kappa, the concentration parameter of a von Mises distribution, showing stronger phase locking during running behavior for CA1 EXC cells (INH mean = 0.41 ± 0.02; EXC mean = 0.63 ± 0.04; ***p = 1.28×10^−10^, Wilcoxon rank sum). Differences in proportions were evaluated using chi-squared tests.

**Fig. S9.**
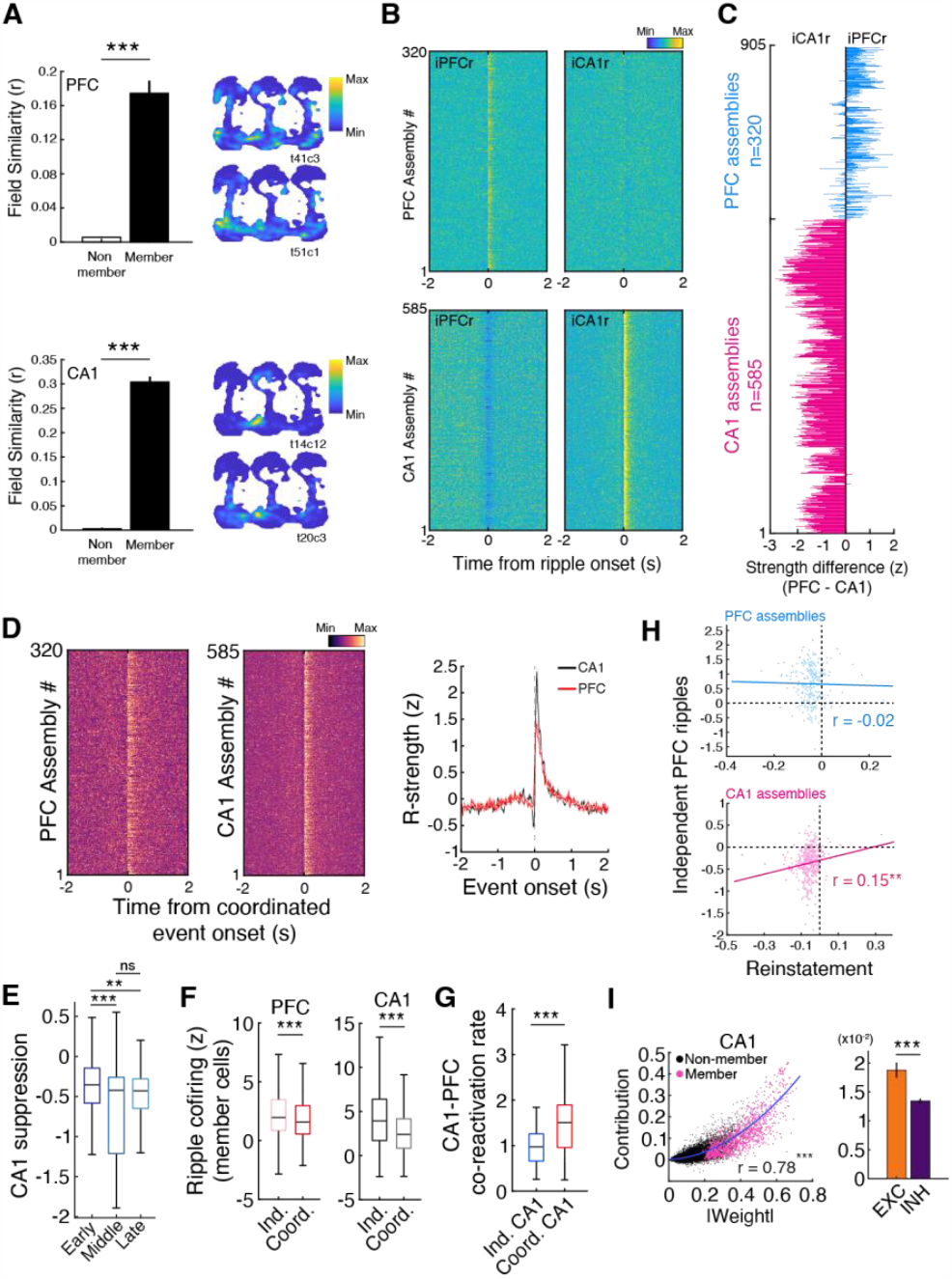
Assembly reactivation in CA1 and PFC. (**A**) Spatial map correlation between member and non-member cells. Field similarity was calculated as the Pearson correlation between the concatenated linearized firing fields of each pair of neurons (CA1: member = 0.30 ± 0.01, non-member = 0.003 ± 2.18×10^−4^, ***p = 1.48×10^−185^; PFC: member = 0.17 ± 0.01, non-member = 0.006 ± 3.96×10^−4^, ***p = 1.75×10^−30^, Wilcoxon rank sum). (**B**) Heat map of all PFC (n = 320) and CA1 (n = 585) assemblies aligned to independent ripple events. (**C**) Assembly reactivation strength difference. This was calculated by taking the mean reactivation strength in a ±200 ms window around independent PFC ripples and subtracting the reactivation strength around independent CA1 SWRs for each assembly. (**D**) Heat map of assemblies aligned to coordinated event onset. The start and end times of a coordinated event were defined as the start and end time of the preceding and following ripple, respectively. (**E**) Suppression of CA1 assemblies during independent PFC ripples grouped by early, middle, and late sleep epochs (Early = -0.39 ± 0.02, Middle = -0.63 ± 0.05, Late = -0.49 ± 0.02, ***p = 2.92×10^−4^, **p = 0.009, p = 0.66 for early vs middle, early vs late, and middle vs late, respectively; Kruskal-Wallis test, Bonferroni corrected for multiple comparison). (**F**) Z-scored ripple co-activity of assembly member cells during independent or coordinated events in their respective area (PFC: Independent = 2.27 ± 0.13, Coordinated = 1.89 ± 0.012, ***p = 8.82×10^−5^; CA1: Independent = 4.72 ± 0.17, Coordinated = 3.09 ± 0.12, ***p = 3.79×10^−67^, Wilcoxon signed-rank). (**G**) CA1-PFC assembly coactivation rate during independent and coordinated ripples (Independent = 1.47 ± 0.20 Hz, Coordinated = 2.60 ± 0.40 Hz, ***p = 5.17×10^−5^, Wilcoxon rank sum). (**H**) Relationship between independent PFC ripple reactivation and reinstatement of assemblies in the subsequent run epoch (PFC: r = - 0.002, p = 0.76, n = 257; CA1: r = 0.15, **p = 0.0031, n = 382). (**I**) Relationship between CA1 assembly weight and contribution (r = 0.78, ***p = 0). CA1 EXC cells had higher overall contribution to assembly strength, consistent with higher absolute weights of EXC cells (Fig. 3D) (EXC = 1.88×10^−2^ ± 1.3×10^−3^, INH = 1.34×10^−2^ ± 4.13×10^−4^, ***p = 5.66×10^−6^, Wilcoxon rank sum).

**Fig. S10.**
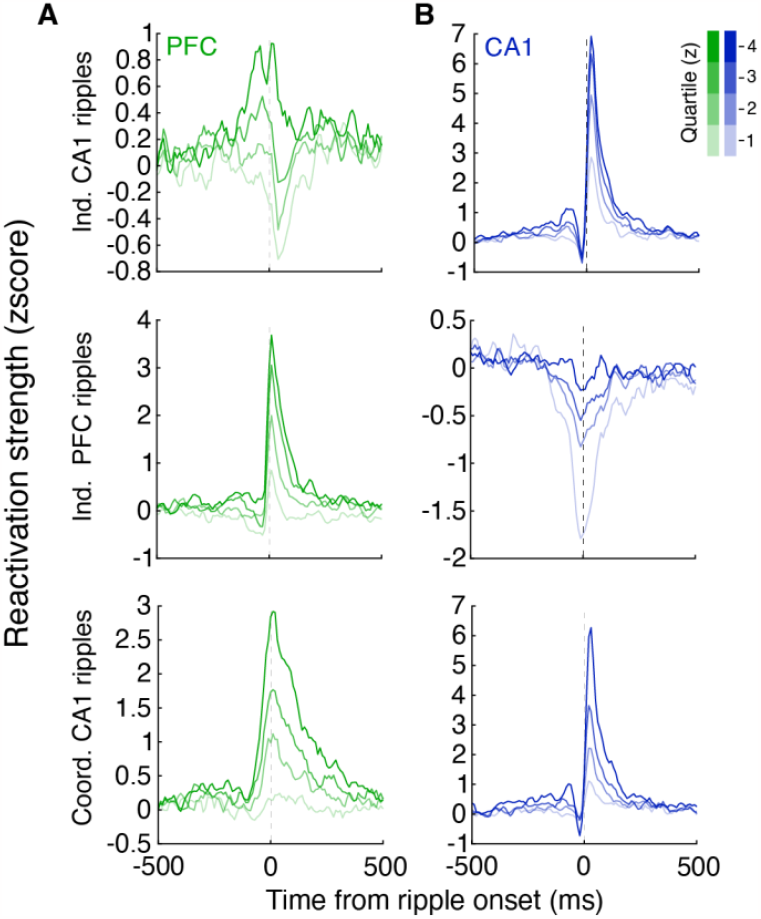
Varying strength of reactivation/suppression across assemblies. (**A**) Reactivation strength of PFC assemblies separated by quartiles to illustrate distinct reactivation profiles across different assemblies. (**B**) Same as (**A**) but for CA1. Assemblies were separated based on their mean reactivation strength in a ±200 ms window around event onset.

**Fig. S11.**
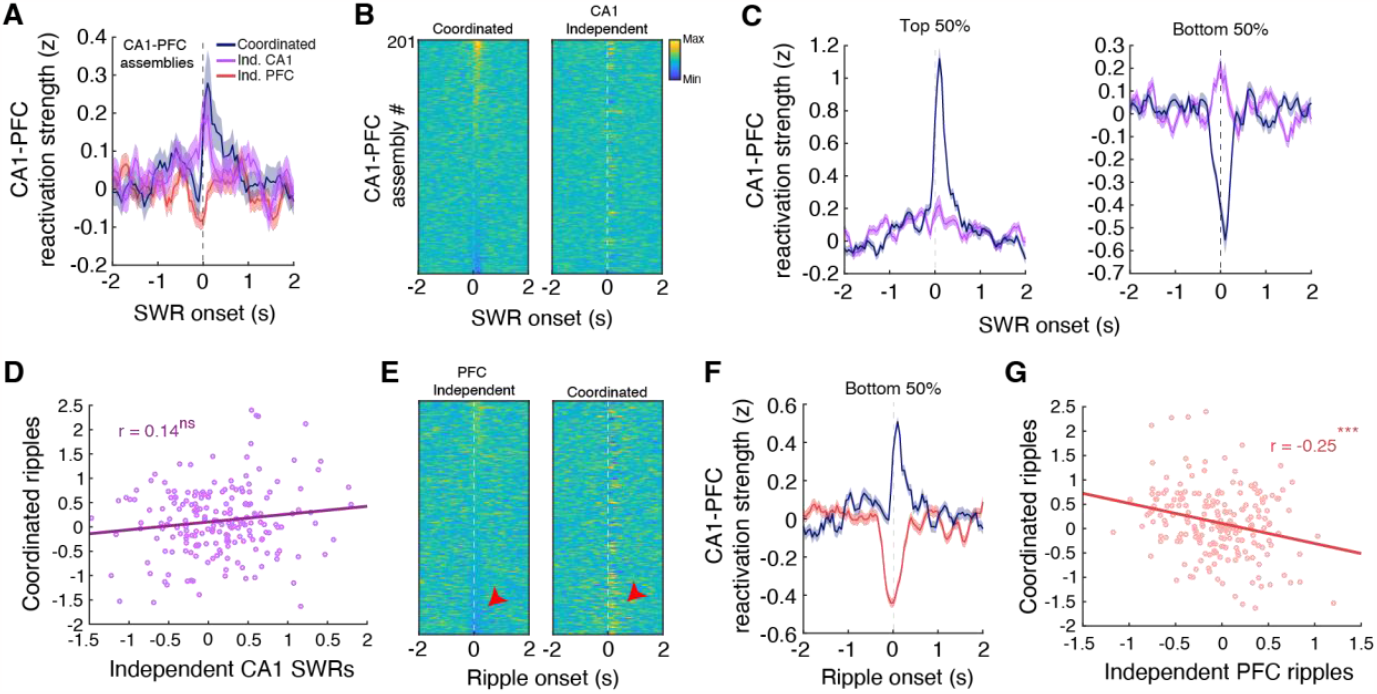
Difference in CA1-PFC coactivity during independent and coordinated ripples. (**A**) Ripple aligned CA1-PFC reactivation strength. (**B**) All assemblies sorted by coordinated SWR aligned reactivation strength. Assemblies must have at least one member cell in each area to be included in analysis. Note that reactivation strength is not consistent across the two SWR types. (**C**) The top and bottom 50% of assemblies sorted by coordinated SWR aligned reactivation strength, further illustrating the difference in coactivity patterns. (**D**) Reactivation strengths during coordinated and independent SWRs were not correlated (r = 0.14, p = 0.052). (**E**) All assemblies sorted by independent PFC ripple aligned reactivation strength. Note that the high degree of CA1-PFC assembly suppression during independent PFC ripples is associated with high reactivation during coordinated ripples (red arrowheads). (**F**) The bottom 50% of assemblies sorted by independent PFC ripple aligned reactivation strength. (**G**) Reactivation strengths during independent PFC ripples were negatively correlated with reactivation during coordinated events, similar to what is seen in CA1 assemblies (r = -0.25, ***p = 3.69×10^−4^).

**Fig. S12.**
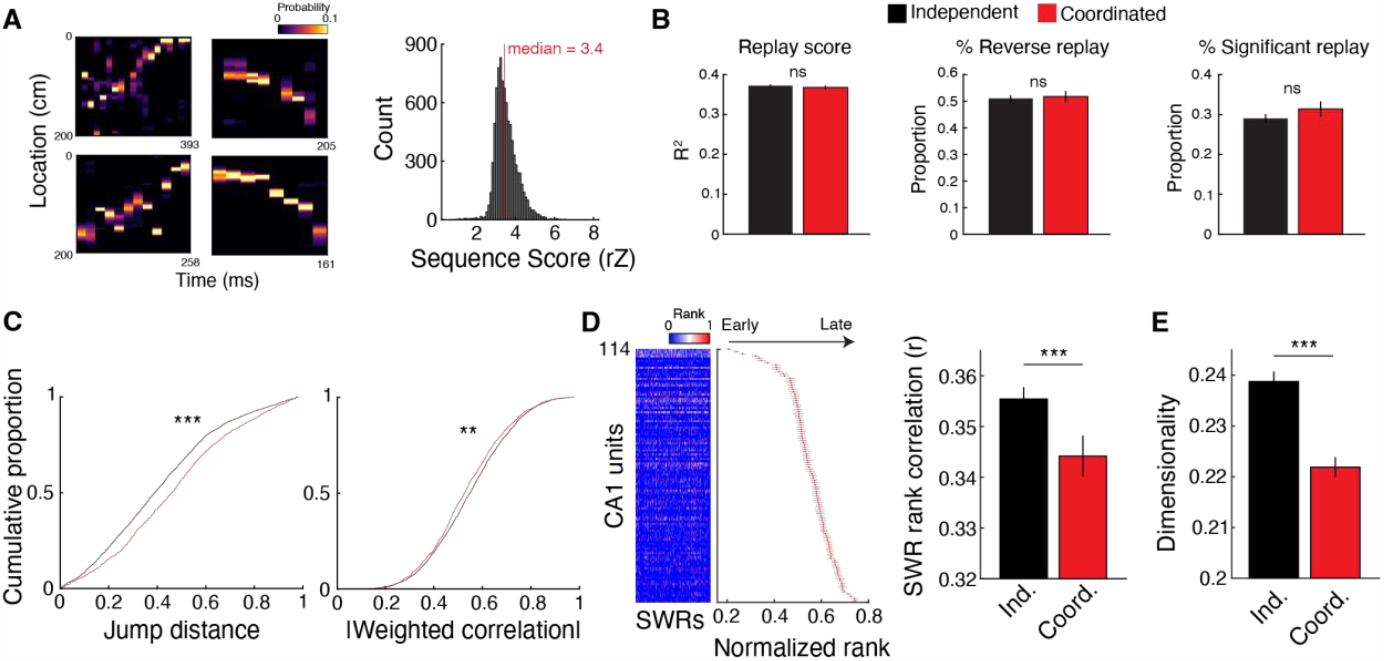
An alternative method of hippocampal replay detection, rank-order analysis, and dimensionality of SWR events. (**A**) Example replay events and sequence scores for each significant event utilizing the weighted correlation shuffling method (median rZ = 3.4). (**B**) Replay score (Independent = 0.372 ± 0.003, Coordinated = 0.368 ± 0.005, p = 0.48, Wilcoxon rank sum), proportion of reverse replay (Independent = 0.51 ± 0.01, Coordinated = 0.52 ± 0.02, p = 0.75, Wilcoxon rank sum), and proportion of significant replay events (Independent = 0.29 ± 0.01, Coordinated = 0.31 ± 0.02, p = 0.28, Wilcoxon rank sum) for independent (n = 3,407) and coordinated (n = 1,357) events assessed by the alternative, line-fitting replay detection method. (**C**) Jump distance (Independent = 0.409 ± 0.004, Coordinated = 0.466 ± 0.007, ***p = 1.34×10^−13^, Wilcoxon rank sum) and weighted correlation (Independent = 0.549 ± 0.003, Coordinated = 0.534 ± 0.004, **p = 0.0014, Wilcoxon rank sum) calculated from significant replay events assessed using the line-fitting method. (**D**) Rank-order correlation analysis. Example rank-order matrix and template from an example sleep epoch (Left). Templates were generated separately for independent and coordinated ripple events. The order of unit spiking (both pyramidal cells and interneurons included) was more consistent for independent SWR events as compared to coordinated SWRs (Independent = 0.356 ± 0.002, Coordinated = 0.344 ± 0.004, ***p = 3.34×10^−5^, Wilcoxon rank sum). (**E**) Population dimensionality was higher during independent SWRs as compared to coordinated SWRs (Independent = 0.239 ± 0.002, Coordinated = 0.222 ± 0.002, ***p = 4.37×10^−15^, Wilcoxon rank sum)

**Fig. S13.**
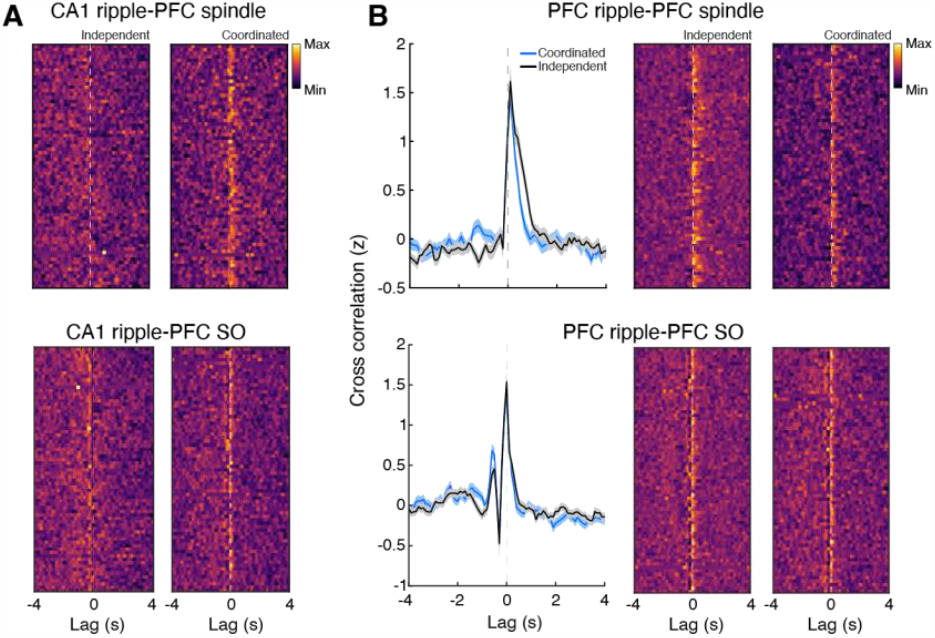
Coupling of slow oscillations and spindles with ripples in CA1 and PFC. (**A**) Heat maps illustrating the cross-correlation between independent or coordinated CA1 SWRs with PFC spindles and SOs (trough time). (**B**) Same as in (**A**), but for PFC ripple events. Each row in the heat map is one epoch from an animal. To calculate the cross-correlation, the ripple time vectors were stationary while the SO or spindle vectors were shifted (lagged). Thus, positive and negative lag values indicate that the oscillatory events tend to precede and follow ripples, respectively.

**Fig. S14.**
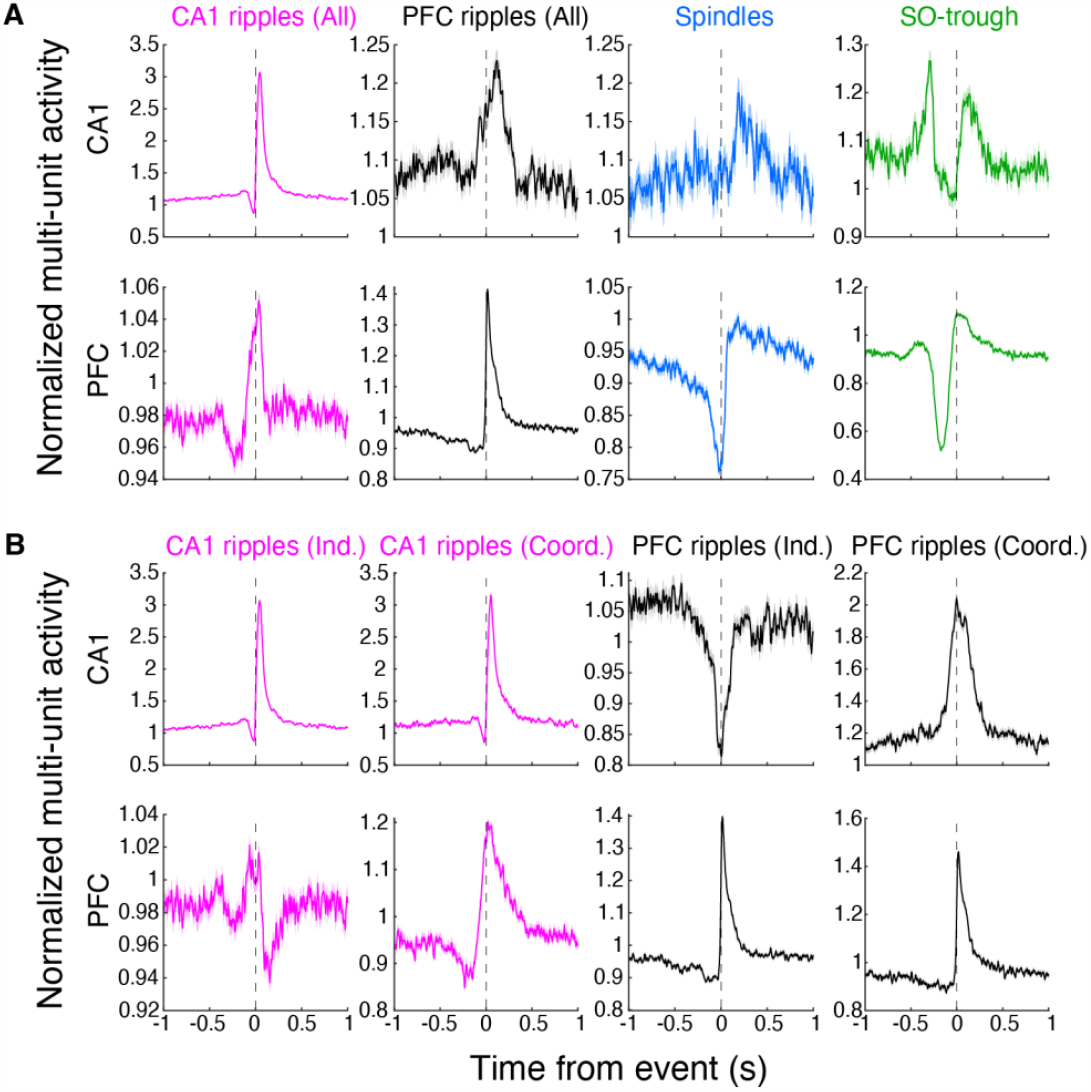
Characterization of population activity surrounding LFP events. (**A**) Multiunit activity in CA1 and PFC aligned to oscillatory events. (**B**) Same is in (**A**) but for independent and coordinated ripples. Population activity was normalized to average activity during NREM sleep.

**Fig. S15.**
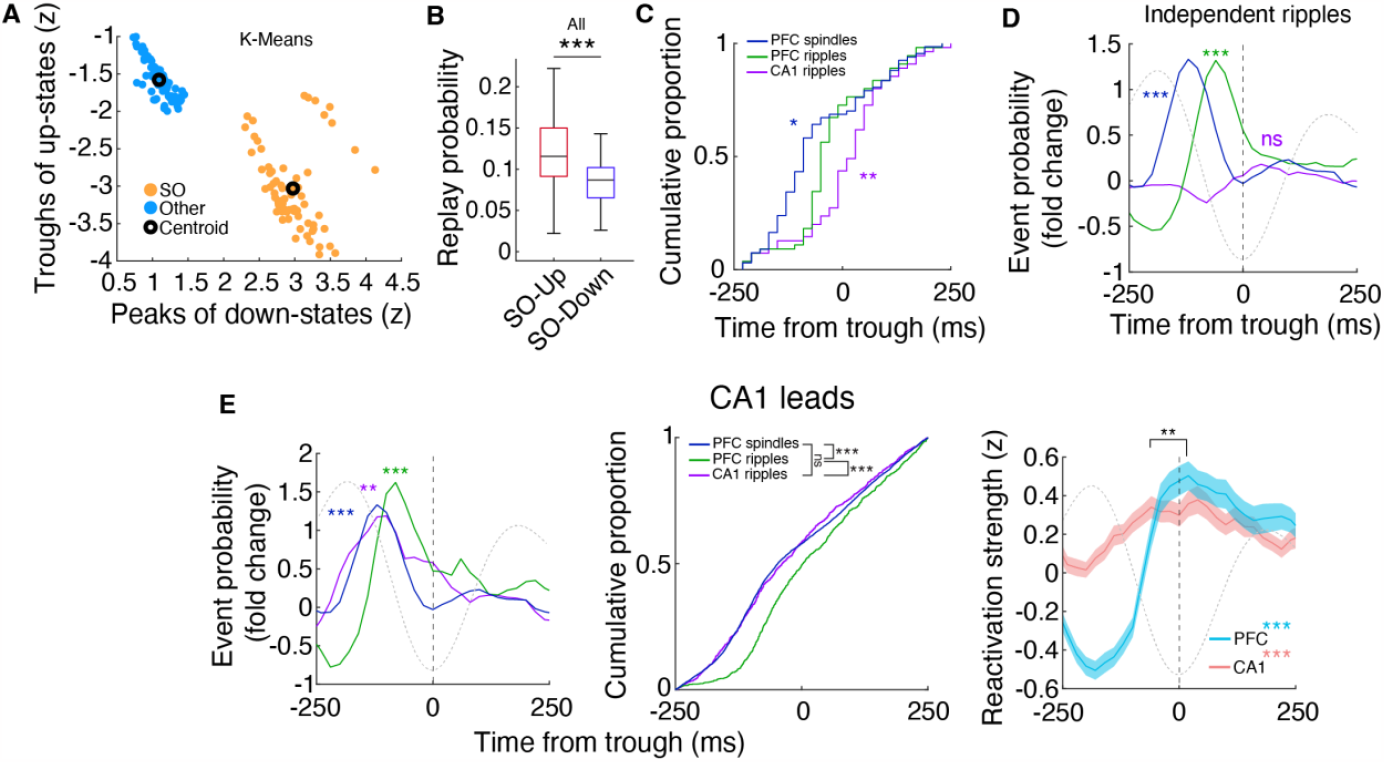
Temporal coordination of events during sleep. (**A**) Designation of high amplitude events as SOs. Note the high degree of separability. Each dot represents the mean peak and trough from an epoch from 1 animal. (**B**) CA1 replay probability during up and down states for all ripples (All: Up = 0.12 ± 0.01, Down = 0.08 ± 0.01, ***p = 7.49×10^−6^, Wilcoxon rank sum). (**C**) Peak timing of spindles, PFC ripples, and CA1 SWRs relative to SO troughs separated by epochs (Spindles vs PFC ripples, *p = 0.01; PFC ripples vs CA1 SWRs, **p = 0.008, Wilcoxon rank sum). (**D**) Event probability fold change for independent PFC and CA1 ripples (Spindle peak, ***p = 2.84×10^−14^; PFC ripple peak, ***p = 4.37×10^−16^; CA1 SWR peak, p = 0.20, ttest vs 0). (**E**). (Left) Event probability fold change for coordinated events where CA1 SWRs precede PFC ripples (Spindle peak, ***p = 2.84×10^−14^; PFC ripple peak, ***p = 4.61×10^−5^; CA1 SWR peak, **p = 0.0019, ttest vs 0). (Middle) Timing of events showed similar distributions for spindles and CA1 SWRs (Spindles vs PFC ripples, ***p = 1.55×10^−18^, Spindles vs CA1 SWRs, p = 0.78, PFC ripples vs CA1 SWRs, ***p = 7.14×10^−12^, Wilcoxon rank sum, Bonferroni corrected). Note that event probability for spindles is the same across all figures since these events were not split into groups (Fig. 4E, fig. S15D,E). (Right) SO trough aligned CA1 and PFC reactivation strength during SO associated coordinated events where CA1 SWRs precede PFC ripples (PFC vs shuffle, ***p = 5.02×10^−6^; CA1 vs shuffle, ***p = 4.32×10^−4^; PFC vs CA1 timing, **p = 0.0041).

**Fig. S16.**
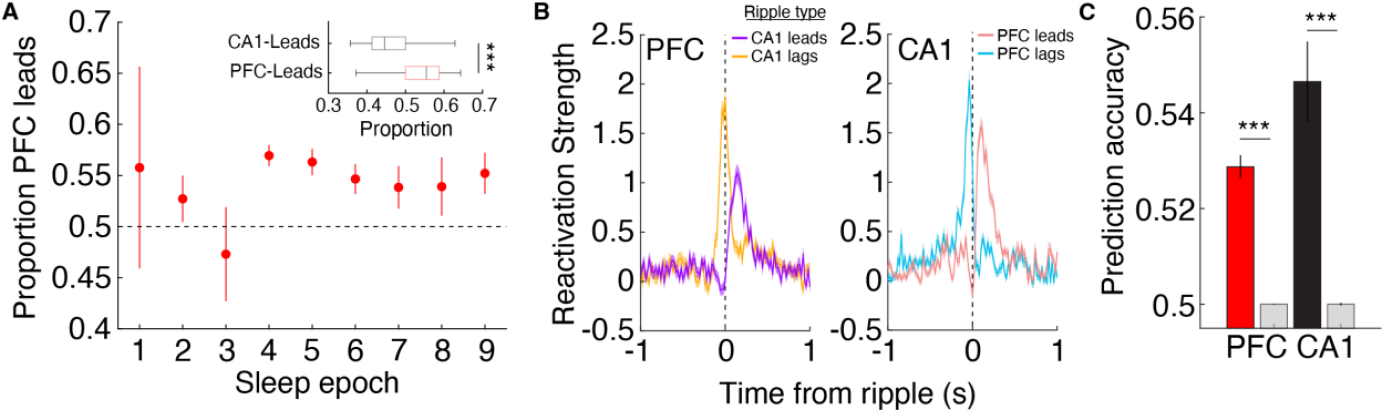
Differences between leading and lagging ripple events. (**A**) PFC ripples tend to precede CA1 ripples during coordinated events (Proportion PFC-leads = 0.54 ± 0.01; ***p = 4.26×10^−9^, Wilcoxon rank sum). (**B**) Comparison of reactivation profiles of PFC and CA1 assemblies aligned to either leading or lagging events in the opposing area illustrate differences in timing of reactivation surrounding these events. (**C**) Joint CA1-PFC spiking activity is able to distinguish leading vs lagging PFC or CA1 ripple events (Predict PFC ripples = 0.53 ± 0.002; shuffle = 0.5 ± 8.1×10^−5^; ***p = 4.87×10^−8^; Predict CA1 ripples = 0.55 ± 0.01; shuffle = 0.5 ± 2.5×10^−4^; ***p = 2.66×10^−18^, Wilcoxon rank sum).

